# Within-sibship GWAS improve estimates of direct genetic effects

**DOI:** 10.1101/2021.03.05.433935

**Authors:** Laurence J Howe, Michel G Nivard, Tim T Morris, Ailin F Hansen, Humaira Rasheed, Yoonsu Cho, Geetha Chittoor, Penelope A Lind, Teemu Palviainen, Matthijs D van der Zee, Rosa Cheesman, Massimo Mangino, Yunzhang Wang, Shuai Li, Lucija Klaric, Scott M Ratliff, Lawrence F Bielak, Marianne Nygaard, Chandra A Reynolds, Jared V Balbona, Christopher R Bauer, Dorret I Boomsma, Aris Baras, Archie Campbell, Harry Campbell, Zhengming Chen, Paraskevi Christofidou, Christina C Dahm, Deepika R Dokuru, Luke M Evans, Eco JC de Geus, Sudheer Giddaluru, Scott D Gordon, K. Paige Harden, Alexandra Havdahl, W. David Hill, Shona M Kerr, Yongkang Kim, Hyeokmoon Kweon, Antti Latvala, Liming Li, Kuang Lin, Pekka Martikainen, Patrik KE Magnusson, Melinda C Mills, Deborah A Lawlor, John D Overton, Nancy L Pedersen, David J Porteous, Jeffrey Reid, Karri Silventoinen, Melissa C Southey, Travis T Mallard, Elliot M Tucker-Drob, Margaret J Wright, Social Science Genetic Association Consortium, Within Family Consortium, John K Hewitt, Matthew C Keller, Michael C Stallings, Kaare Christensen, Sharon LR Kardia, Patricia A Peyser, Jennifer A Smith, James F Wilson, John L Hopper, Sara Hägg, Tim D Spector, Jean-Baptiste Pingault, Robert Plomin, Meike Bartels, Nicholas G Martin, Anne E Justice, Iona Y Millwood, Kristian Hveem, Øyvind Naess, Cristen J Willer, Bjørn Olav Åsvold, Philipp D Koellinger, Jaakko Kaprio, Sarah E Medland, Robin G Walters, Daniel J Benjamin, Patrick Turley, David M Evans, George Davey Smith, Caroline Hayward, Ben Brumpton, Gibran Hemani, Neil M Davies

**Author notes:** these authors contributed equally (Ben Brumpton, Gibran Hemani, Neil M Davies).

## Abstract

Estimates from genome-wide association studies (GWAS) represent a combination of the effect of inherited genetic variation (direct effects), demography (population stratification, assortative mating) and genetic nurture from relatives (indirect genetic effects). GWAS using family-based designs can control for demography and indirect genetic effects, but large-scale family datasets have been lacking. We combined data on 159,701 siblings from 17 cohorts to generate population (between-family) and within-sibship (within-family) estimates of genome-wide genetic associations for 25 phenotypes. We demonstrate that existing GWAS associations for height, educational attainment, smoking, depressive symptoms, age at first birth and cognitive ability overestimate direct effects. We show that estimates of SNP-heritability, genetic correlations and Mendelian randomization involving these phenotypes substantially differ when calculated using within-sibship estimates. For example, genetic correlations between educational attainment and height largely disappear. In contrast, analyses of most clinical phenotypes (e.g. LDL-cholesterol) were generally consistent between population and within-sibship models. We also report compelling evidence of polygenic adaptation on taller human height using within-sibship data. Large-scale family datasets provide new opportunities to quantify direct effects of genetic variation on human traits and diseases.

## Main

Genome-wide association studies (GWAS) have identified thousands of genetic variants associated with complex phenotypes [1, 2], typically using samples of non-closely related individuals [3]. GWAS associations can be interpreted as estimates of direct individual genetic effects, i.e., the effect of inheriting a genetic variant (or a correlated variant) [4–6]. However, there is growing evidence that GWAS associations estimated from samples of unrelated individuals also capture effects of demography [7, 8] (assortative mating [9–11], population stratification [12]) and indirect genetic effects of relatives [13–19] **(Figure 1: panel A).** These non-direct sources of genetic associations are themselves of interest (e.g., for estimating parental effects [13,18], understanding human mate choice [9–11] and genomic prediction [14,19]) but could bias downstream analyses using GWAS summary data such as biological annotation, heritability estimation [20–22], genetic correlations [23], Mendelian randomization [7, 24, 25] and polygenic adaptation tests [26–28].

**Figure 1A.**
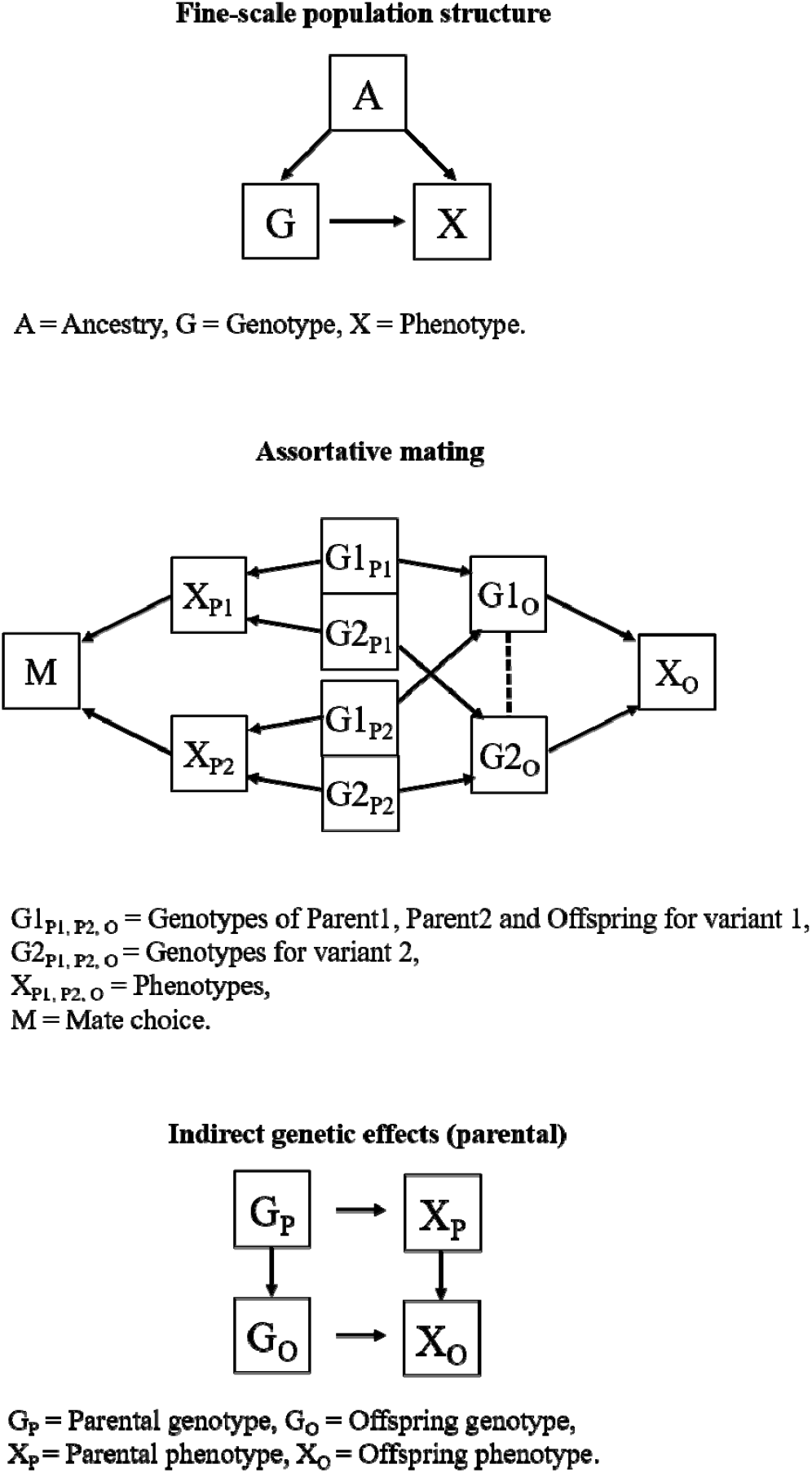
Demographic and indirect genetic effects

**Figure 1B.**
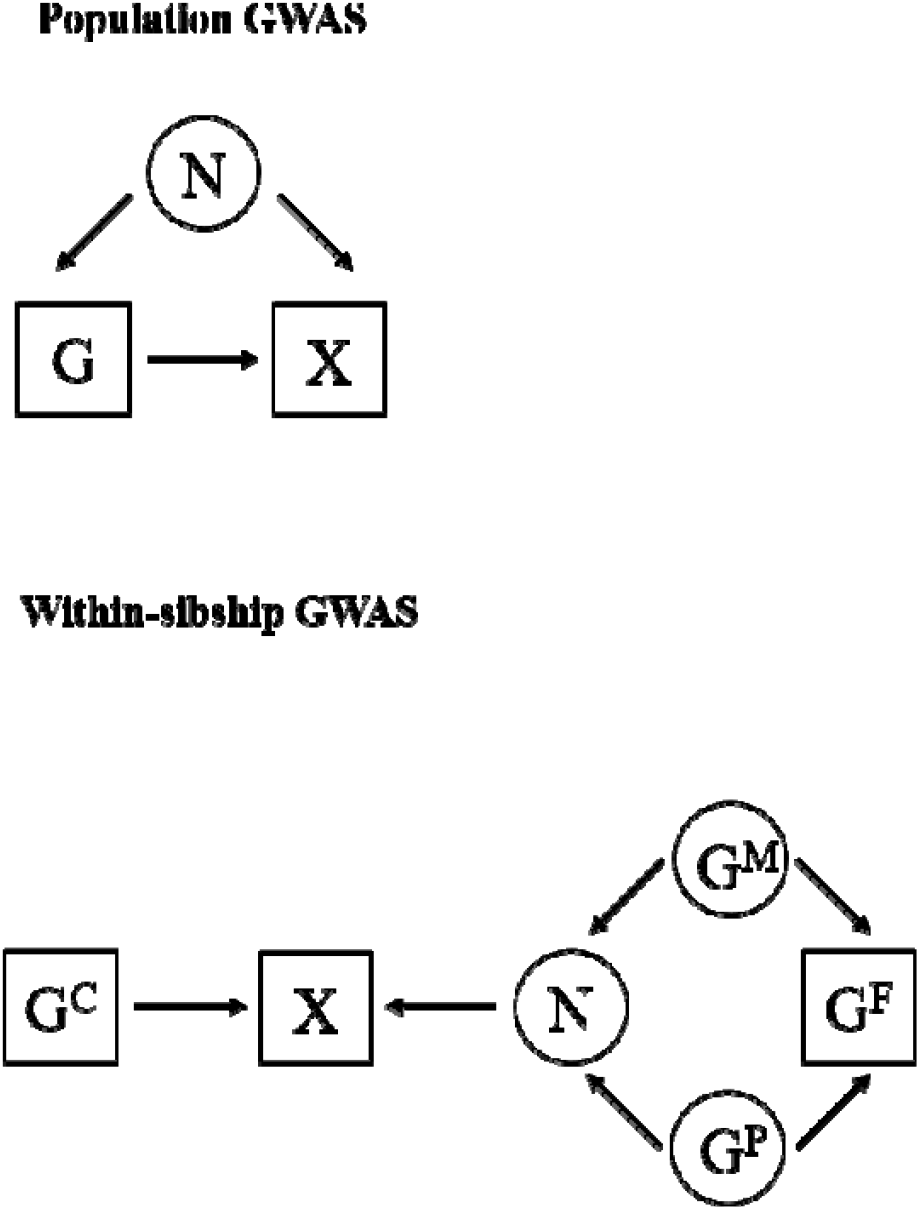
Population and within-sibship GWAS

Within-family genetic association estimates, such as those obtained from samples of siblings, can provide more accurate estimates of direct genetic effects because they are unaffected by demography and indirect genetic effects of parents [7, 17, 29–32]. GWAS using siblings (within-sibship GWAS) **(Figure 1: panel B)** have been previously limited by available data, but are now feasible by combining well-established family studies with recent large biobanks that incidentally or by design contain thousands of sibships [33–36].

Here, we report findings from a within-sibship GWAS of 25 phenotypes using up to 159,701 siblings from 17 studies, the largest GWAS conducted within-sibships to date **(Figure 1: panel C).** We demonstrate that population GWAS estimates for at least 6 phenotypes (height, educational attainment, age at first birth, cognitive ability, depressive symptoms and smoking) partially reflect demography and indirect genetic effects, which affect downstream analyses such as estimates of heritability and genetic correlations. However, we find that associations with clinical phenotypes, such as lipids, are less likely to be affected. We found strong evidence of polygenic adaption on taller human height using within-sibship data. Our study illustrates the importance of collecting GWAS data from families for understanding the effects of inherited genetic variation, particularly for phenotypes sensitive to assortative mating, population stratification and indirect genetic effects.

**Figure 1C.**
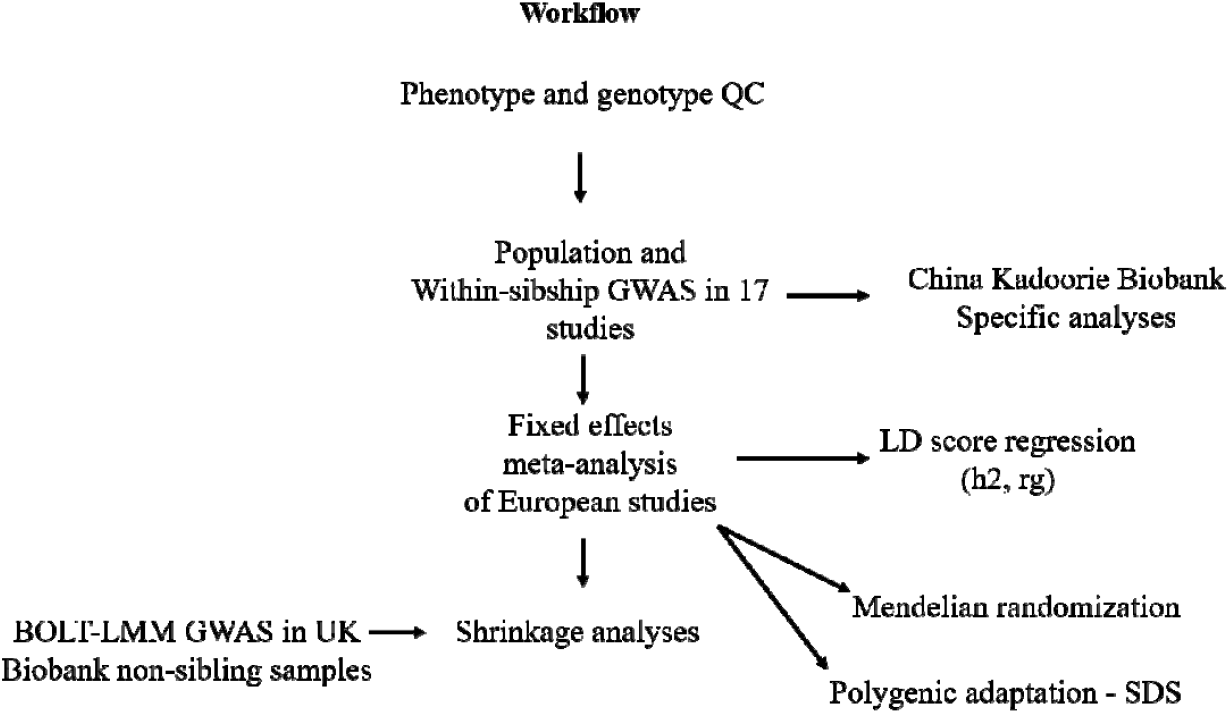
Flowchart illustrating datasets and analyses

## Results

### Genetic association estimates differ when accounting for demography and indirect genetic effects

For GWAS analyses we used 159,701 individuals (with one or more genotyped siblings) from 68,691 sibships in 17 studies **(Supplementary Table 1**). We used within-sibship models which, instead of estimating genotypic associations using the individual’s raw genotypes and population-based samples of unrelated individuals, uses deviations of the individual’s genotype from the mean genotype within the sibship (i.e. all siblings in the family present in the study). Here, the within-sibship model includes the mean sibling genotype as a covariate to capture the between-family contribution of the SNP [14]. For comparison, we also applied a standard population GWAS model that does not account for mean sibship in the same samples. Standard errors were clustered by sibship for both models. All analyses were performed in individual studies and meta-analyses were conducted across studies using summary data. Amongst the phenotypes analysed, the largest available sample sizes were for height (N = 152,350), body mass index (BMI) (N = 144,757), ever smoking (N = 125,949), systolic blood pressure (SBP) (N = 123,406) and educational attainment (N =123,084) **(Supplementary Table 2),** we also report stratified results from non-European samples.

Previous studies have found that association estimates of height and educational attainment genetic variants are smaller in within-family models [13, 14, 37]. We aimed to investigate whether similar shrinkage is observed for other phenotypes by comparing within-sibship and population genetic association estimates for 25 phenotypes. We observed the largest within-sibship shrinkage for genetic variants associated with age at first birth (49%, 95% C.I. [25%, 74%]) and educational attainment (46%, [40%, 52%]). We also found evidence of shrinkage for depressive symptoms (39%, [5%, 73%]), ever smoking (19%, [9%, 30%]), cognitive ability (18%, [2%, 35%]) and height (10%, [8%, 12%]). In contrast, within-sibship association estimates for C-reactive protein (CRP) were larger than population estimates (−9%, [−15%, −2%]). We found limited evidence of within-sibship differences for the remaining 18 phenotypes, including BMI and SBP **(Figure 2 / Supplementary Table 3).**

**Figure 2.**
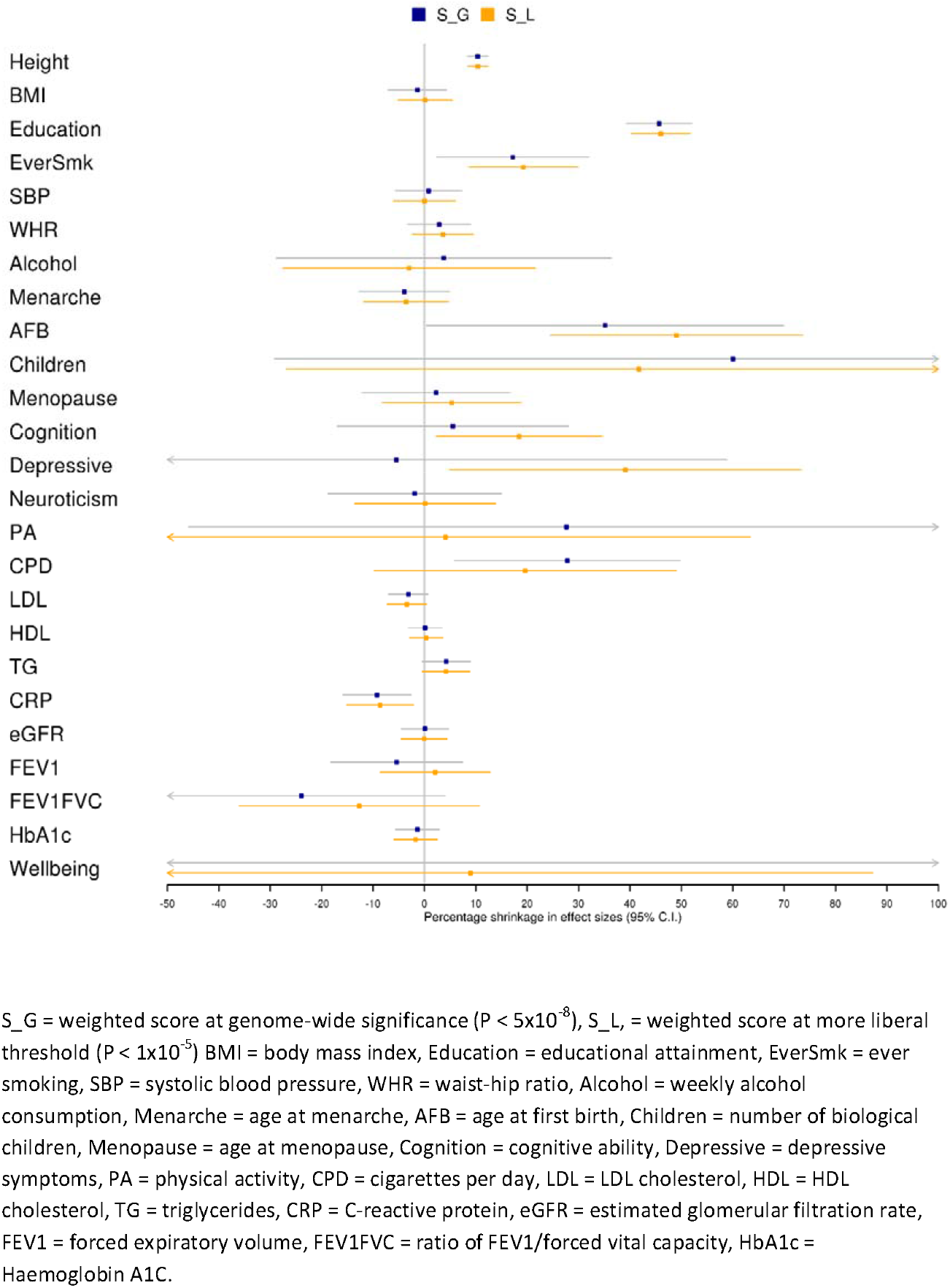
Effect size shrinkage in within-sibship models

We investigated possible heterogeneity in shrinkage for height and educational attainment genetic variants across individual variants and between cohorts. In the meta-analysis data, we observed minimal evidence of heterogeneity in shrinkage across individual variants for height and educational attainment, suggesting that shrinkage is largely uniform across the strongest association signals for these phenotypes. We also found limited evidence of cohort heterogeneity in shrinkages for height (heterogeneity P = 0.25) and educational attainment (P = 0.17) across the European-ancestry cohorts **(Supplementary Figures 1/2).** In contrast, there was limited evidence for shrinkage on height in China Kadoorie Biobank (shrinkage −3%; 95% C.I. [−13%, 7%]; heterogeneity with European meta-analysis P-value = 0.005) but some evidence of shrinkage on ever smoking (shrinkage = 134%; [10%, 258%]) **(Supplementary Figure 3).**

### Within-sibship SNP heritability estimates

LD score regression (LDSC) can use GWAS data to estimate SNP heritability, the proportion of phenotypic variation explained by common SNPs [20, 23]. We used simulations to investigate the applicability of LDSC when using within-sibship GWAS data, finding evidence that LDSC can estimate SNP heritability using both population and within-sibship model GWAS data if effective sample sizes (based on standard errors) are used to account for differences in power between the models **(Methods).**

To evaluate the impact of controlling for demography and indirect genetic effects, we compared LDSC SNP heritability estimates based on population and within-sibship effect estimates for 25 phenotypes. Theoretically, within-sibship shrinkage in GWAS effect estimates will also lead to attenuations in within-sibship SNP heritability estimates **(Methods).** The within-sibship SNP heritability point estimate for educational attainment attenuated by 71% from the population estimate (Population h^2^: 0.14, within-sibship h^2^: 0.04, difference P = 1.5 × 10^−20^) with attenuations also observed for cognition (Population h^2^: 0.24, within-sibship h^2^: 0.13, attenuation 46%; difference P = 0.012) and height (Population h^2^: 0.41, within-sibship h^2^: 0.34, attenuation 17%; difference P = 1.3 × 10^−3^). The observed attenuations were consistent with theoretical expectation **(Supplementary Table 4),** suggesting that the lower within-sibship SNP heritability estimates are explained by association estimate shrinkage. Across the 22 additional phenotypes, population and within-sibship SNP heritability estimates were relatively consistent **(Figure 3 / Supplementary Table 5).**

**Figure 3.**
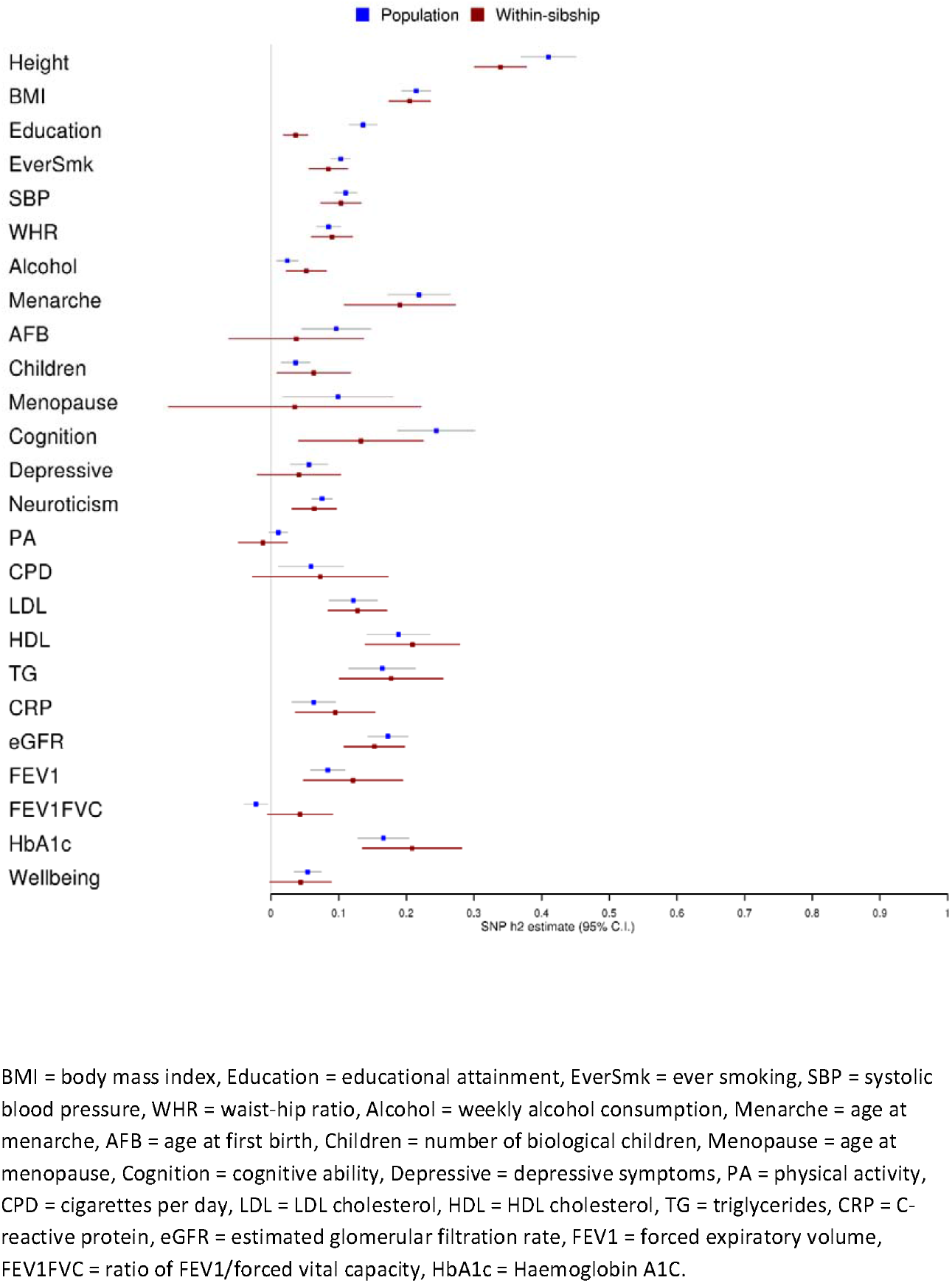
Observed-scale LDSC SNP heritability estimates for 25 phenotypes

### Within-sibship genetic correlations with educational attainment

We used LDSC [23] to estimate cross-phenotype genome-wide genetic correlations (*r_g_*) between educational attainment and 22 phenotypes with sufficient heritability (Population/within-sibship h^2^ > 0). To determine the effects of demography and indirect genetic effects on *r_g_*, we compared estimates of r_g_ using population and within-sibship estimates.

There was strong evidence using population estimates that educational attainment is positively correlated with height (r_g_ = 0.16, 95% C.I [0.10, 0.22]) and negatively correlated with BMI (r_g_ = −0.32, [-0.38, −0.26]) and CRP (r_g_ = −0.47, [−0.69, −0.25]). However, these correlations were negligible when using within-sibship estimates; height (r_g_ = −0.02, [−0.15, 0.10]), BMI (r_g_ = 0.01, [−0.17, 0.19]) and CRP (r_g_ = 0.04, [−0.34,0.42]) with evidence for differences between population and within-sibship r_g_ estimates (height difference P = 4.0×10^−3^, BMI difference P = 6.9×10^−5^, CRP difference P = 0.012). These attenuations indicate that genetic correlations between educational attainment and these phenotypes from population estimates are likely to be driven by demography and indirect genetic effects **(Figure 4 / Supplementary Table 6).**

**Figure 4.**
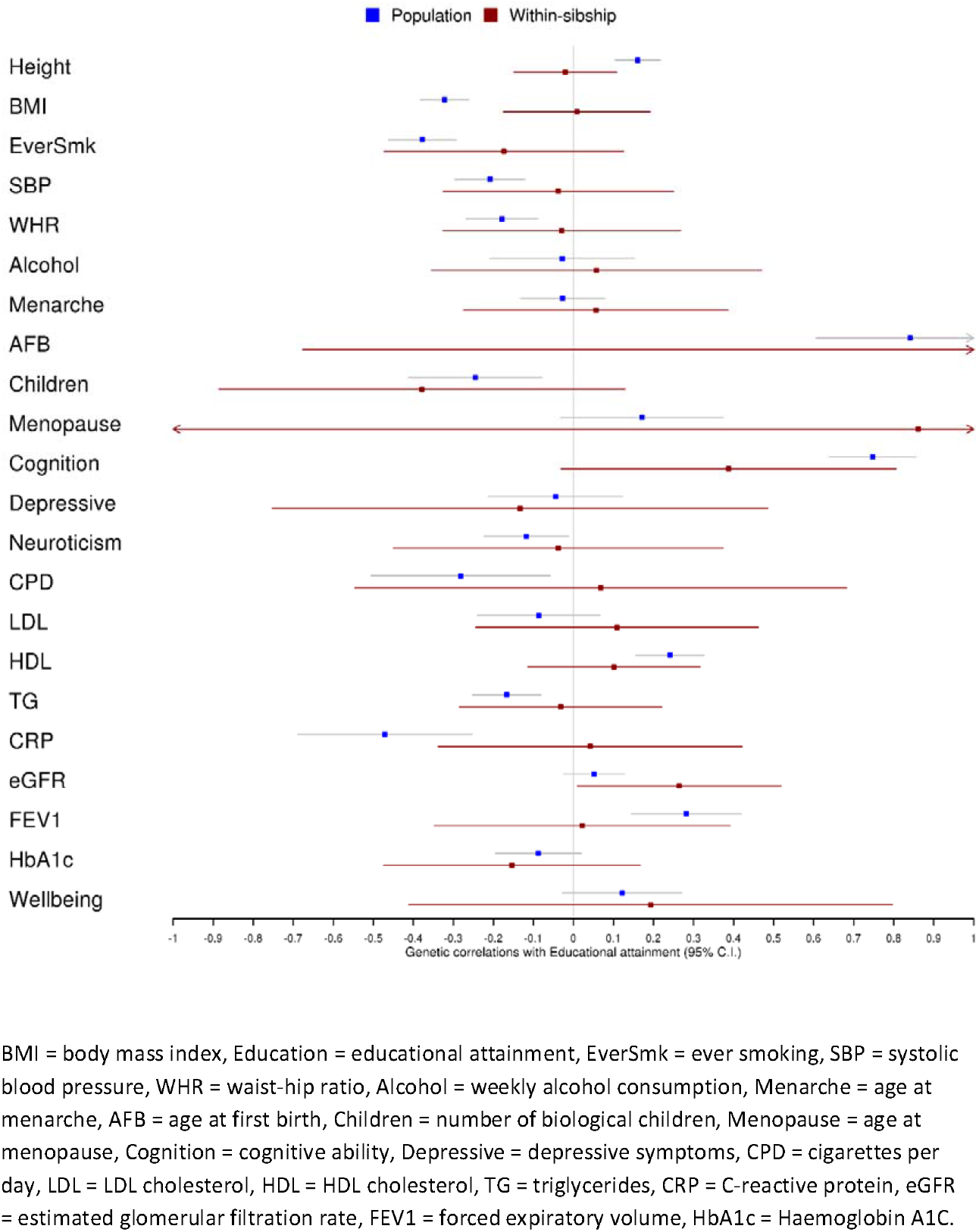
LDSC genetic correlations of phenotypes with educational attainment

### Within-sibship Mendelian randomization: effects of height and BMI

Mendelian randomization uses genetic variants as instrumental variables to assess the causal effect of exposure phenotypes on outcomes [24, 38]. Mendelian randomization was originally conceptualised in the context of parent-offspring trios where offspring inherit a random allele from each parent [24]. However, with limited family data, most Mendelian randomization studies have used data from unrelated individuals. Within-sibship Mendelian randomization (WS-MR) is largely robust against demography and indirect genetic effects that could distort estimates from non-family designs [7, 25]. Here, we used population and WS-MR to estimate the effects of height and BMI on 23 phenotypes.

WS-MR estimates for height and BMI on the 23 outcome phenotypes were largely consistent with population MR estimates for height (0%; 95% C.I. −12%, 12%) but slightly lower for BMI (−11%; 95% C.I. −20%, −1%). However, consistent with the genetic correlation analyses, we observed differences between population and WS-MR estimates of height and BMI on educational attainment. Population Mendelian randomization estimates provided strong evidence that taller height and lower BMI increase educational attainment (0.05 SD increase in education per SD taller height, 95% C.I. [0.03, 0.06]; 0.18 SD decrease in education per SD higher BMI, [0.14, 0.21]). In contrast, WS-MR estimates for these relationships were greatly attenuated (height: 0.01 SD increase [−0.01, 0.03], difference P = 2.9×10^−3^ | BMI; 0.04 SD decrease [0.00, 0.08], difference P = 2.1×10^−7^). We also observed similar attenuation from population and WS-MR estimates for BMI on age at first birth (difference P = 6.0×10^−4^); a phenotype highly correlated with education, and some suggestive evidence of attenuation in the WS-MR estimate of height on triglycerides (difference P = 0.04). These differences illustrate instances where population based Mendelian randomization estimates are distorted by demography and indirect genetic effects **(Table 1**).

**Table 1.**
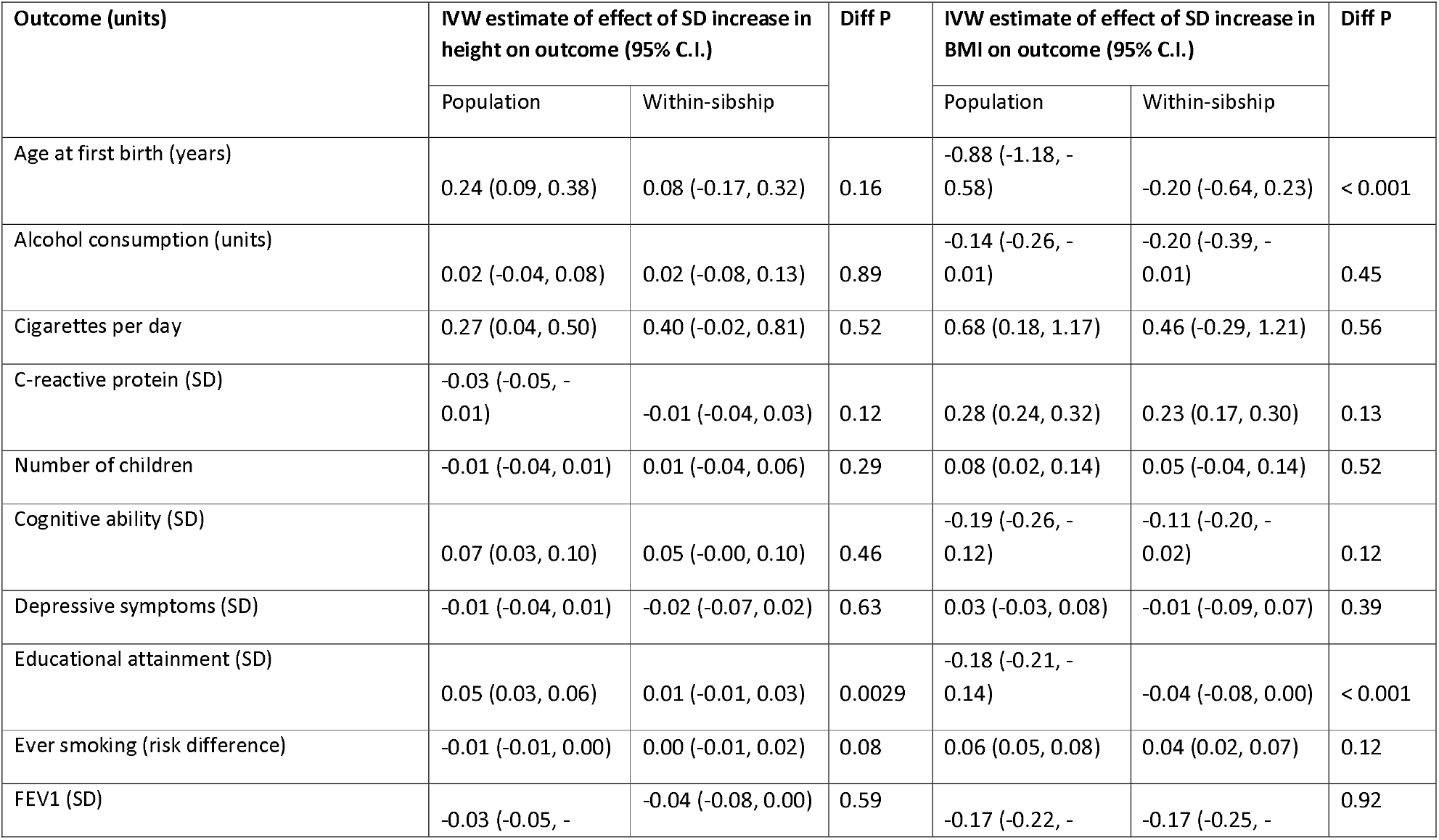
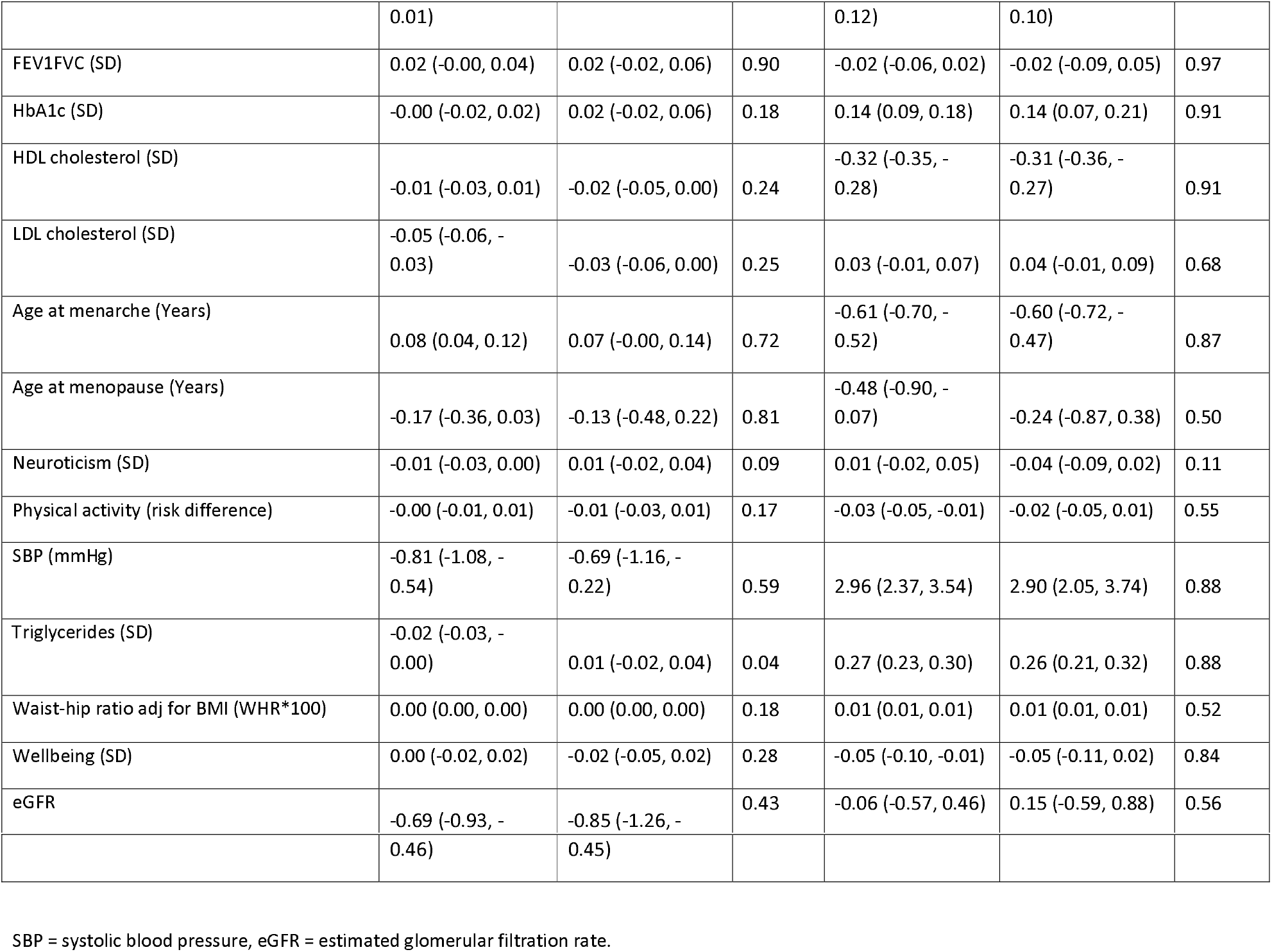
Within-sibship Mendelian randomization: effects of height and BMI on 23 phenotypes

| Outcome (units) | IVW estimate of effect of SD increase in height on outcome (95% C.I.) |  | Diff P | IVW estimate of effect of SD increase in BMI on outcome (95% C.I.) |  | Diff P |
| --- | --- | --- | --- | --- | --- | --- |
|  | Population | Within-sibship |  | Population | Within-sibship |  |
| Age at first birth (years) | 0.24 (0.09, 0.38) | 0.08 (-0.17, 0.32) | 0.16 | -0.88 (-1.18, -0.58) | -0.20 (-0.64, 0.23) | < 0.001 |
| Alcohol consumption (units) | 0.02 (-0.04, 0.08) | 0.02 (-0.08, 0.13) | 0.89 | -0.14 (-0.26, -0.01) | -0.20 (-0.39, -0.01) | 0.45 |
| Cigarettes per day | 0.27 (0.04, 0.50) | 0.40 (-0.02, 0.81) | 0.52 | 0.68 (0.18, 1.17) | 0.46 (-0.29, 1.21) | 0.56 |
| C-reactive protein (SD) | -0.03 (-0.05, -0.01) | -0.01 (-0.04, 0.03) | 0.12 | 0.28 (0.24, 0.32) | 0.23 (0.17, 0.30) | 0.13 |
| Number of children | -0.01 (-0.04, 0.01) | 0.01 (-0.04, 0.06) | 0.29 | 0.08 (0.02, 0.14) | 0.05 (-0.04, 0.14) | 0.52 |
| Cognitive ability (SD) | 0.07 (0.03, 0.10) | 0.05 (-0.00, 0.10) | 0.46 | -0.19 (-0.26, -0.12) | -0.11 (-0.20, -0.02) | 0.12 |
| Depressive symptoms (SD) | -0.01 (-0.04, 0.01) | -0.02 (-0.07, 0.02) | 0.63 | 0.03 (-0.03, 0.08) | -0.01 (-0.09, 0.07) | 0.39 |
| Educational attainment (SD) | 0.05 (0.03, 0.06) | 0.01 (-0.01, 0.03) | 0.0029 | -0.18 (-0.21, -0.14) | -0.04 (-0.08, 0.00) | < 0.001 |
| Ever smoking (risk difference) | -0.01 (-0.01, 0.00) | 0.00 (-0.01, 0.02) | 0.08 | 0.06 (0.05, 0.08) | 0.04 (0.02, 0.07) | 0.12 |
| FEV1 (SD) | -0.03 (-0.05, - | -0.04 (-0.08, 0.00) | 0.59 | -0.17 (-0.22, - | -0.17 (-0.25, - | 0.92 |
|  | 0.01) |  |  | 0.12) | 0.10) |  |
| FEV1FVC (SD) | 0.02 (-0.00, 0.04) | 0.02 (-0.02, 0.06) | 0.90 | -0.02 (-0.06, 0.02) | -0.02 (-0.09, 0.05) | 0.97 |
| HbA1c (SD) | -0.00 (-0.02, 0.02) | 0.02 (-0.02, 0.06) | 0.18 | 0.14 (0.09, 0.18) | 0.14 (0.07, 0.21) | 0.91 |
| HDL cholesterol (SD) | -0.01 (-0.03, 0.01) | -0.02 (-0.05, 0.00) | 0.24 | -0.32 (-0.35, -0.28) | -0.31 (-0.36, -0.27) | 0.91 |
| LDL cholesterol (SD) | -0.05 (-0.06, -0.03) | -0.03 (-0.06, 0.00) | 0.25 | 0.03 (-0.01, 0.07) | 0.04 (-0.01, 0.09) | 0.68 |
| Age at menarche (Years) | 0.08 (0.04, 0.12) | 0.07 (-0.00, 0.14) | 0.72 | -0.61 (-0.70, -0.52) | -0.60 (-0.72, -0.47) | 0.87 |
| Age at menopause (Years) | -0.17 (-0.36, 0.03) | -0.13 (-0.48, 0.22) | 0.81 | -0.48 (-0.90, -0.07) | -0.24 (-0.87, 0.38) | 0.50 |
| Neuroticism (SD) | -0.01 (-0.03, 0.00) | 0.01 (-0.02, 0.04) | 0.09 | 0.01 (-0.02, 0.05) | -0.04 (-0.09, 0.02) | 0.11 |
| Physical activity (risk difference) | -0.00 (-0.01, 0.01) | -0.01 (-0.03, 0.01) | 0.17 | -0.03 (-0.05, -0.01) | -0.02 (-0.05, 0.01) | 0.55 |
| SBP (mmHg) | -0.81 (-1.08, -0.54) | -0.69 (-1.16, -0.22) | 0.59 | 2.96 (2.37, 3.54) | 2.90 (2.05, 3.74) | 0.88 |
| Triglycerides (SD) | -0.02 (-0.03, -0.00) | 0.01 (-0.02, 0.04) | 0.04 | 0.27 (0.23, 0.30) | 0.26 (0.21, 0.32) | 0.88 |
| Waist-hip ratio adj for BMI (WHR*100) | 0.00 (0.00, 0.00) | 0.00 (0.00, 0.00) | 0.18 | 0.01 (0.01, 0.01) | 0.01 (0.01, 0.01) | 0.52 |
| Wellbeing (SD) | 0.00 (-0.02, 0.02) | -0.02 (-0.05, 0.02) | 0.28 | -0.05 (-0.10, -0.01) | -0.05 (-0.11, 0.02) | 0.84 |
| eGFR | -0.69 (-0.93, - | -0.85 (-1.26, - | 0.43 | -0.06 (-0.57, 0.46) | 0.15 (-0.59, 0.88) | 0.56 |
|  | 0.46) | 0.45) |  |  |  |  |

SBP = systolic blood pressure, eGFR = estimated glomerular filtration rate.

### Polygenic adaptation

Polygenic adaptation is a process via which phenotypic changes in a population over time are induced by small shifts in allele frequencies across thousands of variants. One method of testing for polygenic adaptation is to compare Singleton Density Scores (SDS), measures of natural selection over the previous 2,000 years [28], with GWAS P-values. However, this approach is sensitive to population stratification as illustrated by recent work using UK Biobank data which showed that population stratification in GWAS data likely confounded previous estimates of polygenic adaptation on height [26, 27]. Within-sibship GWAS data is particularly useful in this context as it is robust against population stratification. Here we re-calculated Spearman’s rank correlation (r) between tSDS (SDS scores aligned with the phenotype increasing allele) and our population/within-sibship GWAS P-values for 25 phenotypes, with standard errors estimated using jack-knifing over blocks of genetic variants.

We found strong evidence for polygenic adaptation on taller height in the European meta-analysis GWAS using both population (r = 0.020, 95% C.I. [0.011, 0.029]) and within-sibship GWAS estimates (r = 0.011, [0.003, 0.020]) **(Supplementary Figures 4/5).** These results were supported by several sensitivity analyses; a) evidence of enrichment for positive tSDS (mean = 0.23, SE = 0.06, P < 0.001) amongst 416 putative height loci from the within-sibship meta-analysis data **(Supplementary Figure 6),** b) positive LDSC *r_g_* between height and tSDS in the meta-analysis data **(Supplementary Table 7)** and c) evidence for polygenic adaptation on taller height when meta-analysing correlation estimates from 7 individual studies (e.g. SDS using only UK Biobank GWAS summary data) for population (r = 0.013, [0.010, 0.015]) and within-sibship (r = 0.004, [0.002, 0.007]) estimates **(Figure 5**). There was also some putative within-sibship evidence for polygenic adaptation on increased number of children (P = 0.006) and lower HDL-cholesterol (P = 0.024) **(Supplementary Figure 4).**

**Figure 5.**
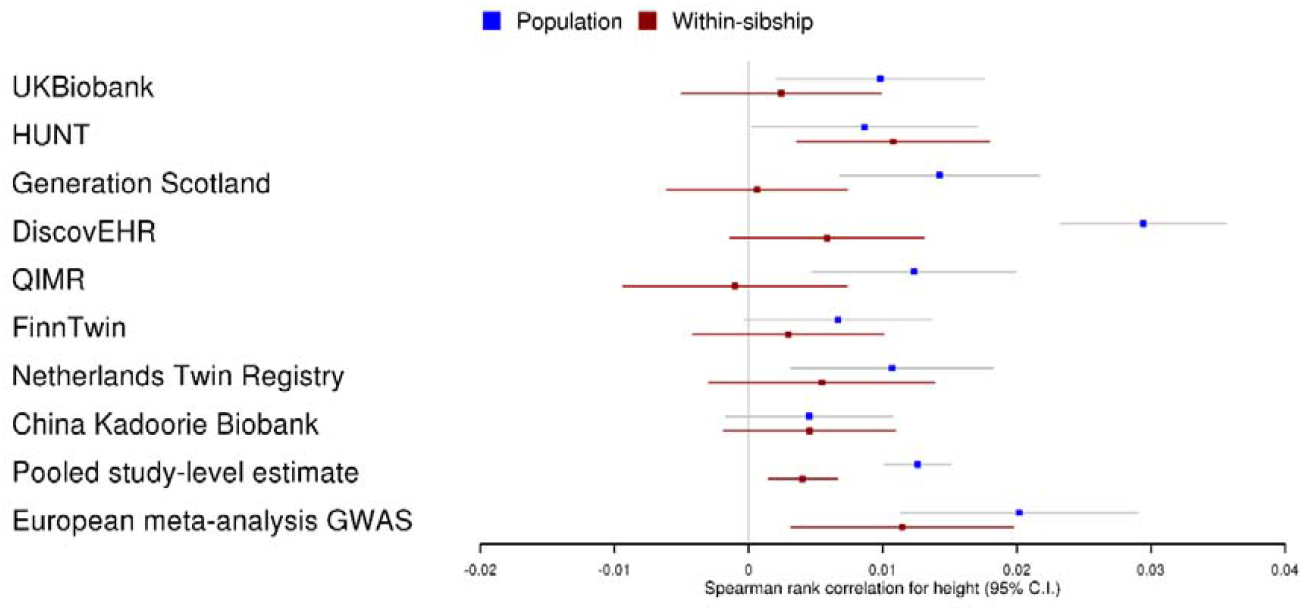
Individual study, pooled and GWAS meta-analysis estimates for polygenic adaptation on height

## Discussion

These results demonstrate that GWAS results and downstream analyses of behavioural phenotypes (e.g. educational attainment, smoking behaviour) as well as some biologically proximal phenotypes (e.g. height, BMI) are likely to be affected by demography and indirect genetic effects. However, we found that most analyses involving more clinical phenotypes, such as lipids, were not strongly affected. Future studies should use data from unrelated individuals, to maximise sample size for gene discovery and polygenic prediction, and data from families to provide more accurate estimates of direct genetic effects for downstream analyses.

A key aim of GWAS is to identify direct genetic effects on phenotypes, but other sources of genetic associations can be of value for analyses. For example, knowledge of indirect genetic effects can be used to elucidate maternal effects [15, 39] or the extent to which health outcomes are mediated by family environments [13, 18]. Future family based GWAS will also enable the estimation of indirect genetic effects [6, 18, 40].

We observed minimal evidence of heterogeneity in shrinkage estimates of height and educational attainment genetic variants. This indicates that observed shrinkage is likely to be largely driven by assortative mating or indirect genetic effects since both of these tend to influence associations proportional to the direct effect (whereas population stratification is likely to have larger effects on ancestrally informative markers). The common environment terms from classical twin studies suggest that there are likely to be indirect genetic effects on educational attainment [41], cognitive ability [42] and smoking [43], but suggest that the observed shrinkage for height is likely to be a consequence of assortative mating [10, 43, 44].

Within-sibship GWAS data can be useful for validating results from larger samples of unrelated individuals. Here, we showed that population and within-sibship Mendelian randomization estimates of height and BMI were generally consistent for 23 outcome phenotypes. However, we observed differences between within-sibship and population Mendelian randomization estimates of height (on educational attainment and triglycerides) and BMI (on educational attainment and age at first birth), suggesting the Mendelian randomization assumptions do not hold for these relationships in samples of unrelated individuals. For future Mendelian randomization studies, within-sibship estimates could elucidate the potential presence of bias [7, 25].

We used non-European data from China Kadoorie Biobank to evaluate whether demography and indirect genetic effects influence GWAS analyses conducted in the Chinese population. In this sample, we found minimal evidence of shrinkage for height genetic variants but (consistent with the European metaanalysis) suggestive evidence of shrinkage for variants associated with smoking initiation. The absence of shrinkage for height suggests that demographic effects such as assortative mating may differ between populations. Larger within-family studies in non-European populations could be used to evaluate population differences in demographic and indirect effects.

We also used the within-sibship GWAS data to evaluate evidence for recent selection. A previous study reporting polygenic adaptation on height in the UK population was found to be biased by population stratification in the GIANT consortium [26–28]. Previous evidence for adaptation on height using siblings in UK Biobank was suggestive of some adaptation, but statistically inconclusive [26]. Here, using within-sibship GWAS estimates from a larger (~4-fold) sample of siblings, we found strong evidence of polygenic adaptation on increased height and some evidence of adaptation on number of children and HDL-cholesterol. We anticipate that future studies on human evolution will benefit from using large within-family datasets such as our resource.

Within-family GWAS are limited by both available family data and statistical inefficiency (homozygosity within-families). To address this issue, future population-based biobanks could recruit the partners, siblings and offspring of study participants. In contrast, conventional GWAS designs sampling unrelated individuals are likely to be the optimal approach to maximise statistical power for discovery GWAS for genetic associations. Indeed, we found that many genotype-phenotype associations from population GWAS models were also observed in within-sibship GWAS, albeit sometimes with attenuated association estimates. A notable limitation of within-sibship models is that they do not control for indirect genetic effects of siblings, i.e. effects of sibling genotypes on the shared environment. Sibling effects have been estimated to be modest compared to parental effects [6, 45] but could have impacted our GWAS estimates. Our findings are also limited to adult phenotypes. Future within-family GWAS (e.g. using parent-offspring trios) could use data from children to evaluate if childhood phenotypes are more strongly affected by indirect genetic effects.

## Methods

### Study participants

Eighteen cohorts contributed data to the overall study **(Supplementary Table 1**). These cohorts were selected on the basis of having at least 500 genotyped siblings (an individual with 1 or more siblings in the study sample) with at least 1 of the 25 phenotypes that were analysed in the study. Detailed information on genotype data, quality control and imputation processes are provided in the **Supplementary Materials.** Individual cohorts defined each phenotype based on suggested definitions from an analysis plan (see the **Supplementary Materials).**

### GWAS analyses

GWAS analyses were performed uniformly across individual studies using automated scripts and a preregistered analysis plan (https://github.com/LaurenceHowe/SiblingGWAS). Scripts checked strand alignment, imputation scores and allele frequencies for the genetic data as well as missingness for covariates and phenotypes. Scripts also summarised covariates and phenotypes and set phenotypes to missing for sibships if only one individual in the sibship has non-missing phenotype data. To harmonise variants for meta-analysis, genetic variants were renamed in a format including information on chromosome, base pair, and polymorphism type (SNP or INDEL). The automated pipeline restricted analyses to common genetic variants (MAF > 0.01) and removed poorly imputed variants (INFO < 0.3). Analyses were restricted to include individuals in a sibship, i.e. a group of two or more full siblings in the study. Monozygotic twins were included if they had an additional sibling in the study.

GWAS analyses involved fitting both population and within-sibship models to the same samples. The population model is synonymous with a conventional principal component adjusted model, and was fit using linear regression in R. The within-sibship model is an extension of the population model including the family mean genotype (the mean genotype of siblings in each family) as a covariate to account for family structure with each individual’s genotype centred around the family mean [7,14]. Age, sex and up to 20 principal components (10 principal components were included in smaller studies at the discretion of study co-authors) were included as covariates in both models.

For individual *j* in sibship *i* with *n_i_* ≥ 2 siblings:

Population model:

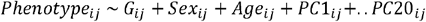

Within-sibship model:

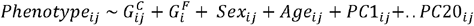

where 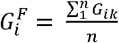 and 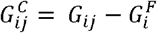

*G_ij_* = genotype of sibling *j* in sibship *i*, 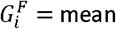 family genotype for sibship *i* over *n* siblings, 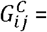 genotype of sibling *j* in sibship *i* centred around 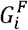.

Standard errors from both estimators were clustered over families at the sibling level to account for nonrandom clustering of siblings within families. Note that this clustering accounts for sibling relationships but does not account for further relatedness present in each sample. For example, a sibling pair could be related to another sibling pair (i.e. two pairs of siblings who are first-cousins). We performed simulations, described below, confirming that such relatedness can lead to underestimating standard errors in the population model and has no effect on the standard errors of the within-sibship model.

GWAS models were performed in individual studies, harmonised and then meta-analysed for each phenotype using a fixed-effects model in METAL [46] with population and within-sibship data meta-analysed separately. We performed meta-analyses using only samples of European ancestry. We used data from 13,856 individuals from China Kadoorie Biobank separately in downstream analyses. The largest meta-analysis sample sizes were for height (N = 137,776), BMI (N = 130,183), ever smoking (N = 111,375), SBP (N = 108,832) and educational attainment (N = 108,510) **(Supplementary Table 2).**

Further quality control was performed prior to meta-analyses. We used phenotype-specific genotype counts (e.g. from the sample with height data in a study) to exclude variants missing in more than 10% of samples. We found some evidence that low frequency variants in small sample sizes may have inflated test statistics in regression models. We randomly selected two sets of 250 sibship (~500 individuals) in UK Biobank and performed population and within-sibship GWAS. We found high levels of test statistic inflation with hundreds (population model) or thousands (within-sibship model) of genome-wide significant hits despite the small sample size. These variants were found to be overwhelmingly low frequency; the 99^th^ percentile MAF for the genome-wide significant variants were 3.0%/2.3% in within-sibship model and 2.9%/4.6% in the population model. Therefore, we used phenotype-specific MAFs and study-level INFO to perform additional stringent quality control on the GWAS data using the following cut-offs for INFO and MAF; fewer than 1,000 individuals (MAF < 0.1, INFO < 0.8); 1,000-3,000 individuals (MAF < 0.05, INFO < 0.5); 3,000-5,000 individuals (MAF < 0.03, INFO < 0.5); 5,000-10,000 individuals (MAF < 0.02, INFO < 0.3); and more than 10,000 individuals (MAF < 0.01, INFO < 0.3).

### Meta-analysis

Phenotypes were harmonised between studies using phenotypic summary data on means and standard deviations. GWAS which did not conform to analysis plan definitions (e.g. binary instead of continuous) were excluded from meta-analyses. GWAS presented in different continuous units (e.g. not standardised) were transformed before meta-analysis by dividing association estimates and standard errors by the standard deviation of the phenotype as measured in the cohort. Meta-analyses for 25 phenotypes were performed using a fixed-effects model in METAL [46].

### Within-sibship and population-based GWAS comparison

#### Overview

We hypothesised that the within-sibship estimates would differ compared to population-based estimates due to the exclusion of effects from demographic and familial pathways. In general, these effects have been shown to inflate (rather than shrink) population-based estimates so we estimated within-sibship shrinkage (the % difference from population to within-sibship estimates). To estimate this shrinkage, we required estimates of the associations with a phenotype from each within-sibship and population-based analyses that were not affected by winner’s curse. Hence, we adopted a strategy where we used an independent reference dataset to select the variants associated with a phenotype. Using the metaanalysis results to obtain association estimates for these variants, we generated summary-based weighted scores of those association estimates in the within-sibship and population-based analyses and estimated the ratio of those scores. We used the UK Biobank dataset that excludes sibling data as the independent reference dataset.

#### GWAS in independent reference discovery dataset

We performed GWAS in an independent sample of UK Biobank (excluding siblings) for each phenotype using a linear mixed model as implemented in BOLT-LMM [47]. We started with a sample of 463,006 individuals of ‘European’ ancestry derived using in-house k-means cluster analysis performed using the first 4 principal components provided by UK Biobank with standard exclusions also removed [48]. To remove sample overlap, we then excluded the sibling sample (N = 40,276), resulting in a final sample of 422,730 individuals. To model population structure in the sample, we used 143,006 directly genotyped SNPs, obtained after filtering on MAF > 0.01; genotyping rate > 0.015; Hardy-Weinberg equilibrium p-value < 0.0001 and LD pruning to an r^2^ threshold of 0.1 using PLINK v2.0 [49]. Age and sex were included in the model as covariates.

All 25 phenotypes (conforming to our phenotype definition) were available in UK Biobank data except for a continuous measure of depressive symptoms. For depressive symptoms, we performed a GWAS of binary depression which was excluded from the meta-analysis (see definition in **Supplementary Materials).** Using the BOLT-LMM UK Biobank GWAS data, we performed strict LD clumping in PLINK v2.0 [49] (r^2^ < 0.001, physical distance threshold = 10,000 kb) to generate independent variants associated with each phenotype at genome-wide significance (P < 5×10^−8^) and at a more liberal threshold (P < 1×10^−5^).

#### Summary-based weighted scores

For a particular phenotype the sets of independent variants obtained from the independent UK Biobank GWAS were used to generate a summary-based weighted score using an inverse variance weighted (IVW) approach [50, 51]:

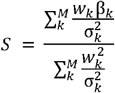

with standard error

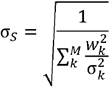

Here the score *S* represents the weighted average of the association estimates of the *M* variants on a phenotype, where β and σ represent the beta coefficients and standard errors from the within-sibship (W) or population-based (P) meta-analysis results. The discovery association estimates from the UK Biobank GWAS were used as weights (w). The set of M variants were determined using either the genome-wide significance (G) or the more liberal threshold (L). Hence, depending on which model is used to determine the association estimates and which set of SNPs are used, for each phenotype four scores can be calculated − *S_P,G_, S_P,L_, S_W,G_* and *S_W,L_*.

These sets of scores were obtained for each of the 25 phenotypes with weights for binary depression used as a substitute for depressive symptoms because a suitable measure was unavailable in UK Biobank. The scores were strongly associated with the set of phenotypes in the meta-analysis data based on determining p-values from their Z-scores. The *S_W,L_* scores were nominally associated at p < 0.05 for 24 out of 25 (exceptions: number of children) of the phenotypes with the *S_P,L_* scores associated with all 25 phenotypes at this threshold **(Supplementary Table 8).**

#### Estimating shrinkage from population to within-sibship estimates

We used the within-sibship and population-based scores to calculate the average shrinkage (δ, i.e. proportion decrease) of genetic variants-phenotype associations

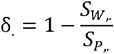

The standard errors of δ could be estimated using the delta method as below using the standard errors of the scores and the covariance between the scores *Cov*(*S_w_, S_P_*):

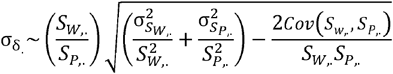

However, we do not have an estimate of this covariance term because the two GWAS were fit in separate regression models. We therefore used the jackknife to estimate σ_δ_. For a score of *M* variants, we removed each variant in turn and repeated IVW and shrinkage analyses as above, extracting the shrinkage point estimate in each of the *M* iterations. We then calculated σ_δ_. as follows:

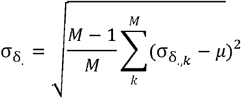

where

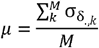

As a sensitivity analysis, we investigated the effects of positive covariance between the population and within-sibship models on the shrinkage standard errors using individual-level participant data from UK Biobank. Analysing shrinkage on height, we used seemingly unrelated regression (SUR) to estimate the covariance term between the population and within-sibship estimators. We found that standard errors for shrinkage estimates decreased by around 15% when the covariance was modelled **(Supplementary Table 9).** SUR standard errors were consistent with the jackknife approach standard errors.

As the primary analysis, we reported shrinkage results using the liberal threshold (P-value < 1×10^−5^) with results using the genome-wide threshold (P-value < 5×10^−8^) reported as a sensitivity analysis. In the main text, we report the shrinkage estimates that reach nominal significance (P < 0.05). We presented shrinkage estimates in terms of % (multiplying by 100).

As a sensitivity analysis, we also presented study-level shrinkage estimates for height and educational attainment and tested for heterogeneity. These phenotypes were chosen because of previous evidence for shrinkage on these phenotypes and available data.

### Heterogeneity of shrinkage across variants within a phenotype

We used results of the within-sibship and population-based meta-analyses to estimate whether shrinkage estimates were consistent across all variants within a phenotype, using an estimate of heterogeneity. As above, we only evaluated heterogeneity for height and educational attainment because of previous evidence and available data. For each variant we estimated the Wald ratio of the shrinkage estimate

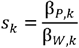

The heterogeneity estimate was obtained as

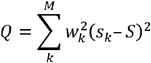

where

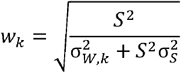

### Applying LD score regression to within-sibship data

LDSC is a widely used method that can be applied to GWAS summary data to estimate heritability and genetic correlation [20, 23]. Central to the method is that LDSC can detect and control for confounding (which is not correlated with LD scores) in GWAS data such as from cryptic relatedness and population stratification. The LDSC ratio, a function of the LDSC intercept unrelated to statistical power, is a measure of the proportion of association signal that is due to confounding. Notably the LDSC ratio will not identify sources of association that are correlated with LD scores such as indirect genetic effects or assortative mating as confounding. We therefore loosely interpret the LDSC ratio as a measure of confounding as it will not identify all sources of confounding. In this work, we apply LDSC to estimate SNP heritability and genetic correlation using the population and within-sibship GWAS data, so we investigated the LDSC intercept/ratio estimates from these data.

In theory, within-sibship data should be less susceptible to confounding than population data as it more effectively controls for population stratification than including principal components. To investigate this in practice, we used LDSC to estimate confounding in meta-analysis summary data for 25 phenotypes. Summary data were harmonised using the LDSC munge_sumstats.py function. LDSC intercepts and ratios were estimated using the harmonised data and the LDSC ldsc.py function with the precomputed European LD scores from the 1000 Genomes (Phase 3) reference panel. The LDSC ratio was used for comparisons between phenotypes and studies as it is not a function of statistical power. The LDSC ratio is calculated from the intercept (i) and the mean chi squared χ^2^as follows:

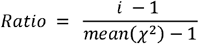

LDSC confounding estimates varied across the 25 phenotypes in the within-sibship model. Confounding estimates were modest for height (11%, 95% C.I. [7%, 14%]) and BMI (6%, [−1%, 14%]) while the estimate for educational attainment was imprecise (35%, [12%, 58%]). Across all phenotypes in the within-sibship data, the median confounding estimate was 21% (Q1-Q3:10%, 33%) but stronger conclusions are limited by imprecise estimates **(Supplementary Table 10/ Supplementary Figure 7).** The LDSC confounding estimates were higher using the population GWAS data (median 44%: Q1-Q3 35%, 56%) than both the within-sibship model and previous studies. For example, the population model LDSC ratio estimates were higher for height (25%, [22%, 28%]), BMI (23%, [20%, 26%]) and educational attainment (47%, [43%, 51%]) **(Supplementary Table 11).**

The observed non-zero confounding in the within-sibship model was unexpected because of the intuition that the within-sibship GWAS models are unlikely to be confounded. The LDSC ratios in the population GWAS were also higher than previous studies. We followed up these findings by evaluating the effects of LD score mismatch and cryptic relatedness on the LDSC ratios.

### Evaluation of LD score mismatch

A large proportion of samples in the meta-analysis were from UK based studies such as UK Biobank and Generation Scotland, for which the LD scores, generated using 1000 Genomes project (phase 3) European samples (CEU, TSC, FIN, GBR), have been shown to fit reasonably well [20]. However, a large number of samples were from Scandinavian populations (HUNT, FinnTwin), where LD mismatch leading to elevated LDSC intercept/ratios has been previously discussed [20]. We investigated this possibility using empirical and simulated data.

We investigated variation in LDSC ratios across populations by comparing ratios for height across well-powered individual studies (N > 5,000): UK Biobank, HUNT, China Kadoorie Biobank (using default East Asian LD scores), Generation Scotland, DiscoverEHR, QIMR and FinnTwin. We found some evidence of heterogeneity between studies; ratio estimates were higher in Scandinavian studies compared to UK-based studies **(Supplementary Figure 8).** We also calculated within-sibship ratio estimates for BMI, SBP and educational attainment using UK Biobank summary data. UK Biobank estimates were largely consistent with zero confounding although confidence intervals were wide **(Supplementary Table 12).**

We performed simulations to evaluate potential mismatch between the Norwegian HUNT study and the default LD scores, which were generated using 1000 Genomes data. We used simulated phenotypes and real genotype data from UK Biobank and HUNT. We estimated the LDSC ratios as above, hypothesising that estimates higher than 0 are likely to reflect LD score mismatch because the phenotypes were simulated to not be influenced by confounders or common environmental terms (which could lead to cryptic relatedness).

Our process was as follows:

a. Select 1,000 HapMap3 SNPs at random.
b. Simulate beta weights for each SNP under a normal distribution with variance defined as a function of allele frequencies. The beta weight for SNP *j* was simulated as follows:
c. *Beta_j_* ~ *N*(0,2*p_j_* (1 – *p_j_*)) where *p_j_* is the minor allele frequency of SNP *j.*
d. Generate polygenic scores for each individual using these weights.
e. Simulate phenotype with 30% of variation explained by polygenic score, with the rest of the variation random.
f. Run GWAS on the simulated phenotype. In UK Biobank we used the Sibling GWAS pipeline on the same sample of siblings. In HUNT we used FastGWA [52] with a sparse GRM on a sample of 30,694 individuals not included in the sibling GWAS sample. The GWAS method and study sample is not particularly important in this context as there were no common environmental effects or confounders in the simulations.
g. Apply LDSC using EUR LD scores to estimate LDSC ratios.

From 10 simulations, the median LDSC ratio estimate was 0.05 (95% C.I. [−0.02, 0.12]) in the population model and 0.05 (95% C.I. [−0.07, 0.16]) in the within-sibship model in UK Biobank, consistent with minimal confounding. In contrast, the median ratio estimate in HUNT was 0.16 (95% C.I. [0.09, 0.23]) when using the default 1000 Genomes LD scores, highly suggestive of non-zero confounding. Using a HUNT-specific LD score reference panel generated using whole genome sequencing data, the median ratio estimate decreased to 0.11 (95% C.I. [0.04, 0.20]) but still suggested non-zero confounding. However, as detailed below, this did not lead to bias in SNP h^2^ estimates.

The combined findings from the empirical and simulated analyses suggest that LD score mismatch with the 1000 Genomes LD scores in HUNT and other studies likely contributed to inflated LDSC ratios in both population and within-sibship GWAS models.

### Cryptic relatedness

One source of inflation in GWAS associations is cryptic relatedness; non-independence between close relatives in the study sample results which leads to inflated precision. In sibling GWAS models we clustered standard errors over sibships, but this clustering does not account for non-independence between related sibships, e.g. uncle/mother and two offspring. Inflated signal relating to cryptic relatedness may result in confounded signal, which is detected by the LD score intercept/ratio. In conventional population GWAS, close relatives are either removed or a mixed model is used to account for close relatives.

The HUNT study population includes many second- and third-degree relatives. To investigate the extent to which cryptic relatedness may have impacted LDSC ratio estimates from the population model, we investigated the effect on the LDSC ratio of using a method that accounts for relatedness. We ran a conventional population GWAS of height using FastGWA [52], which accounts for close relatives using a sparse GRM (IBD > 0.05). We included age, sex, batch and the first 20 principal components as covariates. Using the GWAS summary data we then estimated the LDSC ratio using the 1000 Genomes reference panel and compared with previously described ratio estimates. We found that the FastGWA LDSC ratio (0.33; 95% C.I. [0.28, 0.39]) was substantially lower than the population model LDSC ratio (0.69; 95% C.I. [0.65, 0.73]) suggesting that cryptic relatedness was a source of inflation in the LDSC ratio for the population model.

Cryptic relatedness is an issue for non-family models but may not be an issue for within-family models. We performed simulations to investigate how cryptic relatedness would affect the standard errors of the population and within-sibship GWAS models.

Simulations included 3 generations (generations 1, 2 and 3), and we considered only a single genetic variant *G. We* assumed random mating across all generations and complete Mendelian inheritance for G. Individuals in generation 1 were all unrelated and after pairing randomly, each pair had 2 offspring (generation 2). Similarly, individuals in generation 2 paired randomly and had 2 offspring (generation 3). Generation 2 contained sibling pairs and Generation 3 contained first cousin quads (i.e. two pairs of siblings who are first cousins).

We simulated a common environmental term *C* for Generation 2, which was identical for the full-siblings. In Generation 3, *C* was defined as the mean of parental *C* in Generation 2. We then simulated a normally distributed phenotype *P* in Generation 3 in which 30% of the variation was explained by *C* and the other 70% of variation was random. Note that *P* is not associated with *G.* We then performed regressions of the genetic variant on the phenotype using the population and within-sibship models, extracting the regression P-values. We repeated these simulations and regressions 10,000 times. We found that the type 1 error rate was inflated in the population model (5.84 %) (i.e. the false positive rate was higher than 5%) but not in the within-sibship model (4.94 %).

These findings suggest that the standard errors in the within-sibship model are not underestimated because of cryptic relatedness relating to common environmental effects shared between relatives. This, cryptic relatedness likely inflated LDSC ratios in the population models but not in the within-sibling data. Code for simulations on cryptic relatedness is available on GitHub (github.com/LaurenceHowe/SiblingGWASPost/blob/master/LDSCsimulations/CrypticRelatednessSims.R).

### Within-sibship SNP heritability estimates

LDSC was used to generate SNP heritability estimates for 25 phenotypes using the LDSC harmonised (see above) meta-analysis summary data. The summary data were harmonised using the LDSC munge_sumstats.py function, and we used the precomputed European LD scores from 1000 Genomes Phase 3.

LDSC requires a sample size parameter N to estimate SNP heritability. For this parameter, we used the effective sample size for each meta-analysis phenotype, equivalent to the number of independent observations. This was estimated as follows using GWAS standard errors, minor allele frequencies and the phenotype standard deviations (after adjusting for covariates).

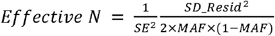

SE = GWAS model standard error, MAF = minor allele frequency of the variant, SD_Resid = standard deviation of the regression residual.

Effective sample size was estimated for each individual study GWAS and each model (e.g. UK Biobank population GWAS of height). To reduce noise from low frequency variants, we restricted to variants with MAF between 0.1 and 0.4 (from 1000 Genomes EUR). At the meta-analysis stage, the effective sample size for each variant was calculated as the sum of sample sizes of studies that the variant was present in.

We used simulated data to validate the use of effective sample sizes and to explore the effects of bias in the LDSC intercept (relating to LD score mismatch) on SNP heritability estimates. In the previously described simulations (in “evaluation of LD score mismatch”) we also estimated SNP heritability alongside the LDSC ratios. In UK Biobank, the median SNP heritability across 10 simulations was 0.29 (95% C.I. [0.23, 0.34]) in the population model and 0.32 (95% C.I. [0.21, 0.42]) in the within-sibship model, highly comparable to the true simulated heritability of 0.30. In HUNT, SNP heritabilities were unbiased using both reference panels, but the median SNP heritability estimate was more precise using the HUNT LD scores (0.31; 95% C.I. [0.25, 0.38]) than the 1000 Genomes LD scores (0.31; 95% C.I. [0.21, 0.42]).

The simulated data suggests that LDSC can generate unbiased estimates of SNP heritability even in the presence of LD score mismatch. However, extensive simulations beyond the scope of this project are required to investigate this further. In empirical analyses, we decided to focus on the differences between the population model 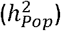 and within-sibship model 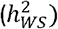 SNP heritability estimates. If we assume that biases affect the estimates equally then the difference between the two estimates will be unbiased. We estimated the difference between the heritability estimates 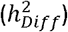 using a difference-of-two-means test [53] as below.

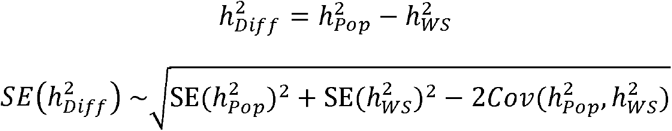

To estimate 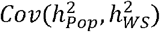, we computed the cross-GWAS LDSC intercept between the population and within-sibship GWAS data (for the same phenotype) which is an estimate of 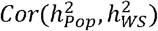. The estimates of this term were ~0.40 across phenotypes. We then calculated the covariance term as follows:

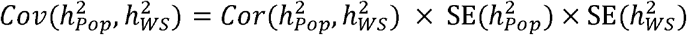

We used the difference Z score (i.e 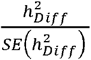) to generate a P-value for the difference between 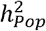 and 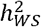. In the text, we report differences reaching nominal significance (difference P < 0.05).

We calculated the expected effect of shrinkage on LDSC SNP heritability estimates. LDSC heritability estimates (*h*^2^) are derived from the formulation below [20]:

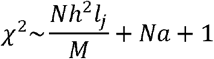

where χ^2^ = the square of the GWAS Z score, *N* = the sample size, *M* = number of variants such that 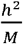 is the average heritability for each variant, *I_j_* is the LD score of variant j, *a* is the effect of confounding biases.

Uniform shrinkage across the genome, would lead to GWAS Z scores being multiplied by a factor (1 – *k*) where *k* is the shrinkage coefficient and *χ^2^* statistics being multiplied by (1 – *k*)^2^. As above, we have used effective sample size to account for differences in *N* between the population and within-sibship models. Therefore, assuming all other coefficients remain consistent,the expectation of 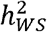 can be written as a function of *k* and 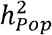.

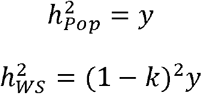

### Within-sibship genetic correlations with educational attainment

We used LDSC to estimate r_g_ between educational attainment and other phenotypes using both population and within-sibship data. LDSC requires non-zero heritability to generate meaningful r_g_ estimates, so we restricted analyses to the 22 phenotypes with SNP heritability point estimates greater than zero in both population and within-sibship models. We estimated the difference between the population (*r_g,Pop_*) and within-sibship (*r_g,WS_*) estimates (*r_g,Diff_*) using a difference-of-two-means test [53],

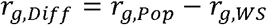

We used the jackknife to estimate the standard error of the difference *SE*(*r_g,Diff_*). After restricting to ~1.2 million Hapmap 3 variants present in the 1000 Genomes LD scores, we ordered variants by chromosome and base-pair and separated variants into 100 blocks. We removed each block in turn and computed *r_g,Diff_* using LDSC 100 times. We then calculated *SE*(*r_g,Diff_*) across the 100 iterations as follows:

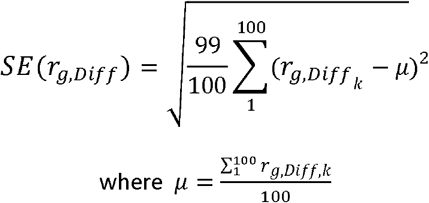

*r_g,Diff,k_* = r_g_ estimate in the kth iteration, *μ* = the mean r_g_ estimate across all 100 iterations We used the difference Z score (i.e. 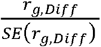) to generate a P-value for heterogeneity between *r_g,Pop_ r_g,WS_*. In the text, we report differences reaching nominal significance (heterogeneity P < 0.05).

### Within-sibship Mendelian randomization: effects of height and BMI

We performed Mendelian randomization analyses using the within-sibship meta-analysis GWAS data to estimate the effect of two exposures (height and BMI) on 23 outcome phenotypes. For the exposure instruments, we used 803 and 418 independent genetic variants for height and BMI, respectively. These variants were identified by LD clumping in PLINK (r^2^ < 0.001, physical distance threshold = 10,000 kb, P < 5×10^−8^) as described in the PRS analysis. We then performed a MR-IVW analysis using the within-sibship meta-analysis data to estimate the effect of the exposure on the outcome as

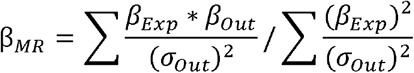

where *β_Exp_* = association estimate from exposure GWAS, *β_Out_* = association estimate from outcome GWAS, *σ_Out_* = standard error from outcome GWAS.

We also performed Mendelian randomization analyses using the population meta-analysis GWAS data for comparison. We estimated differences between population and within-sibship Mendelian randomization estimates using the difference-of-two-means test [53]:

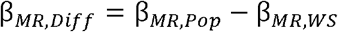

We used the jackknife to estimate the standard error of the difference *SE*(β_*MR,Diff*_) With *n* genetic instruments, we removed each variant from the analysis in turn and then computed β_*MR,Diff*_ storing the estimate from the *n* iterations. We then calculated β_*MR,Diff*_ as follows:

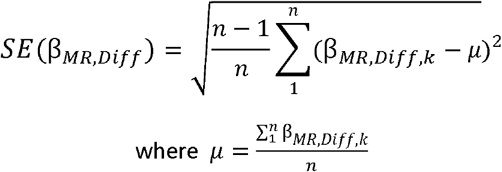

*n* = number of genetic variants used as instruments, β_*MR,Diff,k*_ = β_*MR,Diff*_ estimate in the kth iteration, *μ* = the mean β_*MR,Diff*_ estimate across all *n* iterations.

We used the difference Z score (i.e. 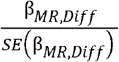) to generate a P-value for heterogeneity between β_*MR,Pop*_ and β_*MR,WS*_. In the text, we report differences reaching nominal significance (heterogeneity P < 0.05).

### Polygenic adaptation

Polygenic adaptation was estimated using similar methods to a previous publication [28]. Precomputed SDS scores were downloaded for UK10K data from https://web.stanford.edu/group/pritchardlab/ Genomic regions under strong recent selection (*MHC* chr6: 25,892,529-33,436,144; lactase chr2: 134,608,646-138,608,646) were removed and SDS scores were normalised within each 1% allele frequency bin.

SDS scores were merged with GWAS meta-analysis data for 25 phenotypes. Variants with low effective sample sizes (< 50% of maximum) were removed for each phenotype. SDS scores were transformed to tSDS such that the reference allele was the phenotype-increasing allele.

Spearman’s rank test was used to estimate the correlation between tSDS and the absolute value of GWAS Z-scores from the population and within-sibship models. Standard errors were estimated using the jackknife. The genome was ordered by chromosome and base pair and divided into 100 blocks. Correlations were estimated 100 times with each kth block removed in turn. The standard error of the correlation estimate *SE*(*Cor*) was calculated as follows:

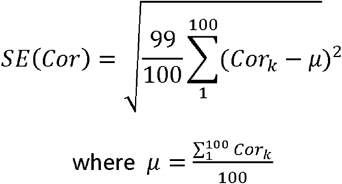

*Cor_k_=* Spearman’s rank correlation estimate in the kth iteration, *μ* = the mean correlation estimate across the 100 iterations.

Given previous concerns [26, 27], we performed several sensitivity analyses for the height analysis. First, we evaluated the mean tSDS in the within-sibship model for a subset of independent variants strongly associated with height. We determined these variants by LD clumping the within-sibship meta-analysis GWAS data in PLINK v 2.0 (r^2^ < 0.001, physical distance threshold = 10,000 kb, P < 1×10^−5^). Second, we used LDSC to estimate the genetic correlation between the SDS scores and the height GWAS data from the population and within-sibship models. The SDS input data was normalised (as above) SDS and we used the precomputed European LD scores from 1000 Genomes. Third, we also calculated spearman rank correlations between height and the SDS (as above) using summary data from individual studies (as opposed to the meta-analysis GWAS). We used all studies with N > 4000, which were UK Biobank, HUNT, Generation Scotland, QIMR, Netherlands Twin Registry, FinnTwin, Discover EHR and China Kadoorie Biobank and investigated both population/within-sibship models. We then used a fixed effects model to meta-analyse the correlation estimates across the studies for the population and within-sibship models. Notably, the correlation estimate using only the UK Biobank WF summary data was inconclusive (r = 0.002; 95% C.I. −0.005, 0.010), consistent with a previous study [26], and correlation point estimates from individual studies were generally smaller than the meta-analysis GWAS estimates. This heterogeneity could relate to the increased number of samples in the meta-analysis, with a higher signal to noise ratio in the individual studies.

## Description for Figure 1A

### Population stratification bias

Population stratification is defined as the distortion of associations between a genotype and a phenotype when ancestry *A* influences both genotype *G* (via differences in allele frequencies) and the phenotype *X*. Principal components and linear mixed model methods control for ancestry but there may be residual confounding relating to fine-scale population structure.

### Assortative mating

Assortative mating is a phenomenon where individuals select a partner based on phenotypic (dis)similarities. For example, tall individuals may prefer a tall partner. Assortative mating can induce correlations between causes of an assorted phenotype in subsequent generations. If a phenotype *X* is influenced by 2 independent genetic variants *G1* and *G2* then assortment on *X* (represented by effects of *X* on mate choice *M*) will induce positive correlations between *G1* in parent 1 and *G2* in parent 2 and vice versa. Parental transmission will then induce correlations between otherwise independent *G1* and *G2* in offspring. These correlations can distort genetic association estimates.

### Indirect genetic effects

Indirect genetic effects are effects of relative genotypes (via relative phenotypes and the shared environment) on the index individual’s phenotype. These indirect effects influence population GWAS estimates because relative genotypes are also associated with genotypes of the index individual. Indirect genetic effects of parents on offspring are of most interest because they are likely to be the largest. However, indirect genetic effects of siblings or more distal relatives are also possible.

## Description for Figure 1B

Population GWAS estimate the association between raw genotypes *G* and phenotypes *X.* As outlined in Figure 1A, estimates from population GWAS may not fully control for effects of demographic biases (population stratification and assortative mating) and may also capture indirect genetic effects of relatives. For simplicity we use *N* to represent all sources of associations between *G* and *X* which do not relate to direct effects of *G.* Circles indicate unmeasured variables and squares indicate measured variables.

If parental genotypes are known, *G* can be separated into non-random (determined by parental genotypes) and random (relating to segregation at meiosis) components.

Within-sibship GWAS include the mean genotype across a sibship (G^F^) (a proxy for the mean of the paternal and maternal genotypes G^P,M^) as a covariate to capture associations between *G* and *X* relating to parents. The within-sibship estimate is defined as the effect of the random component; i.e. the association between family-mean centred genotype G^C^ (i.e. G – G^F^) and *X.* Demographic

Figure 2 displays estimates of shrinkage between population and within-sibship models. Shrinkage is defined as the % decrease in association between the relevant weighted score and phenotype when comparing the population estimate to the within-sibship estimate.

Figure 3 displays LDSC SNP h^2^ estimates for 25 phenotypes using population and within-sibship meta-analysis data.

Figure 4 displays LDSC rg estimates between educational attainment and 22 phenotypes using population and within-sibship meta-analysis data.

Figure 5 displays spearman rank correlation estimates between tSDS (SDS scores aligned with height increasing alleles) and absolute height Z scores. Positive correlations indicate evidence of historical positive selection on height increasing alleles. The pooled estimate is a meta-analysis of the correlation estimates from the individual studies shown above while the European meta-analysis estimate is the correlation estimate using the metaanalysis GWAS data.

Table 1 contains population and within-sibship Mendelian randomization estimates of height and BMI on 23 phenotypes. Units are presented in terms of a standard deviation increase in height or BMI. Difference (diff) P-values refer to evidence of differences between population and within-sibship estimates.

## Acknowledgements

We would like to thank Hakhamanesh Mostafavi and Jonathan Pritchard for helpful suggestions and guidance relating to the polygenic adaptation analyses.

## Contributions

LJH, MGN, TTM, YC, JBP, JFW, JLH, SL, MCS, DAL, NGM, AH, KH, CJW, BOÅ, PDK, JK, SEM, DJB, PT, DME, GDS, CH, BMB, GH and NMD were closely involved in conceptualising and designing the study.

JK, KPH, EMTD, SMK, HC, JFW, EJCD, RP, JAS, PAP, SLRK, SL, JLH, MCS, KC, NMD, SEM, NGM, BMB, RGW, IYM, KL, KH, CJW, CRB, AEJ, DP, CH, AC were involved in data and funding acquisition.

LJH developed the GitHub GWAS pipeline with support from GH and NMD and programming code from GH and PT (via SSGAC). CH kindly beta tested the GWAS pipeline and suggested improvements. Other analysts (listed below) also made major contributions to the GWAS pipeline.

LJH, SG, AFH, HR, CH, YC, GC, PAL, TP, MVDZ, RC, MM, YW, SL, LK, SMR, LFB, CAR, MN, JVB performed GWAS analyses in individual cohorts with the support and guidance of NMD, GH, BMB, RGW, IYM, KL, AEJ, SEM, JK, MGN, MB, JBP, SH, JLH, JFW, JAS, PAP, SLRK, KC, MCK.

LJH performed meta-analyses and all downstream analyses with the meta-analysis data.

LJH drafted the first version of the manuscript. MGN, TTM BOÅ, PDK, JK, SEM, RGW, DJB, PT, DME, GDS, CH, BMB, GH and NMD played a key role in interpreting the results, planning additional analyses, and revising the manuscript.

All authors contributed to and critically reviewed the manuscript.

## Data availability

GWAS summary statistics will be made publicly available for download prior to publication.

