## Supplemental information and figures for "Within-sibship GWAS improve estimates of direct genetic effects"

Howe et al 2021

[Supplementary Figure 6 Histogram of tSDS for independent variants associated with height in the within-sibship meta-analysis data (P < 1x10^-5^)	35](#_Toc65612842)

### Cohort and consortium information

#### Members of the Social Science Genetic Association Consortium (SSGAC)

*Daniel J. Benjamin*, David Cesarini, *Philipp Koellinger*, *Patrick Turley*, Aysu Okbay, Alexander I. Young, Michelle Meyer, Jonathan Beauchamp, Richard Karlsson Linnér, Fleur Meddens, Cornelius A. Rietveld, Ronald de Vlaming, Raymond Walters, Casper Burik, *Hermon Kweon*, Samantha Cherney, Chelsea Watson, Michael Bennett, Tami Gjorgjieva, Grant Goldman, Hariharan Jayashankar, Seyed Moeen Nehzati, Nancy Wang.

Italics denotes co-authorship on the manuscript.

#### Members of the Within-Family Consortium (WFC)

Michel Boivin, Tom A Bond, Laura D Howe, Amanda Hughes, William G Iacono, James J Lee, Marcus R Munafò, Michael C Neale, Juan R Ordoñana, Eleanor Sanderson, Fartein A Torvik, Nicole M Warrington.

#### Cohort descriptions

##### Australian Mammographic Density Twins and Sisters Study

###### Overview

Australian Mammographic Density Twins and Sisters Study (AMDTSS) is a twin and family study for studying mammographic density (Odefrey et al 2010). Female twins and their sisters without breast cancer were recruited between 2004 and 2009. Participants completed questionnaire surveys through telephone-administered interviews to be collected for self-reported weight, height, and other known and putative breast cancer risk factors. Blood samples were collected, couriered to the laboratory within 48 hours of collection, and were processed to generate dried blood spot Guthrie cards. The study was approved by the Human Research Ethics Committee of The University of Melbourne, and written informed consent was obtained from the participants. 1811 first-degree sisters (monozygotic twin pairs were excluded if there was no other sisters available) of 685 sibships were included in this study.

###### Genotyping, imputation and quality control

Genome-wide SNP data were genotyped using DNA extracted from blood samples and the Infinium OncoArray-500K Beadchip. Pre-imputation quality control included filtering SNPs to have call rate >=95%, MAF>=0.01 and HWE P>=10-7, and filtering samples to have call rate>0.95, sex consistency and heterozygosity rate Z<4.89. Imputation was conducted using the Michigan Imputation Server with HRC Release 1.1 as the reference panel. Post-imputation quality control included filtering SNPs to have MAF>=0.01, imputation quality info>=0.3, genotype rate >=95% and HWE P=10-7.

###### Acknowledgements

We would like to thank all participants of the AMDTSS. The AMDTSS was facilitated through access to Twins Research Australia, a national resource supported by a Centre of Research Excellence Grant (number 1079102) from the NHMRC. The AMDTSS was supported by NHMRC (grant numbers 1050561 and 1079102), Cancer Australia and National Breast Cancer Foundation (grant number 509307).

##### The Center on Antisocial Drug Dependence (CADD)

###### Overview

The Center on Antisocial Drug Dependence (CADD) was established in 1997 to investigate the etiology of substance use disorders and related conditions. This study includes individuals from two cohorts within CADD: the longitudinal twin sample (LTS) and the community twin sample (CTS) (Rhea et al., 2013). Recruited in assistance with the Colorado Department of Health (CDH), the LTS sample consists of twins recruited at birth and tested annually. The CTS sample, in comparison, was recruited in collaboration with both CDH and the Colorado Department of Education to include twins ranging in age from 12-18 at the first point of assessment. In both samples, the closest-in-age sibling was included. All analyses were performed using Wave 2 of the LTS and CTS data.

###### Genotyping, imputation and quality control

Individuals were genotyped using the Affymetrix Axiom Precision Medicine Research Array (PMRA) Chip. Pre-imputation quality control included filtering SNPs to have call rate ≥ 95%, MAF ≥ 0.0001, HWE *p* ≥ 10-6, and filtering samples to have call rate > 0.95. Imputation was conducted using EAGLE (version 2.4). Post-imputation quality control included filtering SNPs to have MAF ≥ 0.0001, imputation quality INFO > 0.3, SNP call rate ≥ 95% and HWE *p* ≥ 10-6.

###### Analysis

Analyses were performed using the supplied scripts and using age, sex, and 20 principal components as covariates. The initial assessment of height and weight in these samples were in inches and pounds. These measures were subsequently converted to metric units to derive height (cm) and BMI (kg/m^2).

###### Acknowledgements

We would like to acknowledge and thank all the participants. This research was supported by the National Institute on Drug Abuse (Grant Numbers: DA038504 and DA011015), and utilized resources from the University of Colorado Boulder Research Computing Group, which is supported by the National Science Foundation (Award Nos. ACI-1532235 and ACI1532236), the University of Colorado Boulder, and Colorado State University.

##### China Kadoorie Biobank (CKB)

###### Overview

China Kadoorie Biobank is a prospective population-based cohort of 512,891 adults aged 30-79 years recruited from 10 geographically defined regions during 2004-2008 (1). The baseline survey collected questionnaire data, physical measurements, and blood samples, including measurement of blood glucose with recording of time since last meal. Five-yearly resurveys are undertaken among a 5% randomly-selected sample of surviving participants, collecting the same information as at the baseline survey plus some additional measures. All participants are followed for cause-specific mortality and morbidity and for any hospital admission, through linkages with registries and health insurance databases. Local, national and international ethics approval was obtained and all participants provided written informed consent.

###### Genotyping, imputation and quality control

Genotyping was conducted using custom Affymetrix Axiom® arrays, with 100,706 unique samples (call rate >0.95, no sex mismatch, no XY aneuploidy, heterozygosity < mean+3SD) and 511,885 variants (call rate >0.98, HWE P>1E-06, batch/plate effect P<1E-06, MAF difference from 1000 genomes EAS <0.2) passing QC. Genotypes were phased using SHAPEIT3 v4.12 and imputed into the 1000 Genomes Phase 3 reference with IMPUTE4 v4.r265. After imputation, variants with MAF<0.005 or info<0.3 were excluded. Genotype probabilities were converted to hard-call genotypes in Plink using a cut-off of 0.499 (i.e. the most probable genotype).

Sib-pairs were initially identified as sample pairs with pi-hat>0.375, excluding parent-child pairs by confirming Z0 >0.05, Z1>0.5. Family structures were checked to ensure all family members were sibs, where necessary excluding individuals to remove putative ½- or ¾-sibling relationships. Sole monozygotic twins (i.e. with no other siblings in the dataset) were excluded.

###### Analysis

Analyses were performed using the supplied scripts, including as covariates: age at baseline; sex, recruitment region, and 12 principal components. Systolic blood pressure was adjusted for use of BP-lowering medication by adding 15 mmHg. Alcohol consumption was derived from questionnaire responses as previously described (2).

###### Acknowledgements

China Kadoorie Biobank was supported as follows: Baseline survey and first re-survey: Hong Kong Kadoorie Charitable Foundation; long-term follow-up: UK Wellcome Trust (212946/Z/18/Z, 202922/Z/16/Z, 104085/Z/14/Z, 088158/Z/09/Z), National Natural Science Foundation of China (91843302), and National Key Research and Development Program of China (2016YFC 0900500, 0900501, 0900504, 1303904). DNA extraction and genotyping: GlaxoSmithKline, UK Medical Research Council (MC_PC_13049, MC-PC-14135). The UK Medical Research Council (MC_UU_00017/1,MC_UU_12026/2 MC_U137686851), Cancer Research UK (C16077/A29186; C500/A16896) and the British Heart Foundation (CH/1996001/9454), provide core funding to the Clinical Trial Service Unit and Epidemiological Studies Unit at Oxford University for the project. CKB acknowledges the study participants and members of the survey teams in each of the 10 regional centres, and the project development and management teams based at Beijing, Oxford and the 10 regional centres.

##### Danish Twin Registry (DTR)

###### Overview

This study included 586 dizygotic twin pairs recruited by the Danish Twin Registry (DTR) as part of the study of Middle-Aged Danish Twins (MADT) and the Longitudinal Study of Aging Danish Twins (LSADT). Briefly, MADT was initiated in 1998 and includes 4,314 twins randomly chosen from the birth years 1931-1952. Surviving participants were revisited from 2008 to 2011 (1), where the blood samples used in this study were collected. LSADT was initiated in 1995 and includes twins aged 70 years and older. Follow-up assessments were conducted every second year through 2005 (1). The individuals included in the present study all participated in the 1997 assessment, where blood samples were collected from same sex twin pairs.

For MADT study participants, data on height, BMI, education, depressive symptoms, age at first birth, and physical activity were collected as part of the 1998 assessment, whereas data on cognition, smoking, alcohol consumption, subjective wellbeing, neuroticism, and number of biological children were collected as part of the 2008-2011 follow-up survey. All data for the LSADT study participants were collected as part of the 1997 assessment.

Written informed consents were obtained from all participants. Collection and use of biological material and survey information were approved by the Regional Scientific Ethical Committees for Southern Denmark, and the study was approved by the Danish Data Protection Agency.

###### Genotyping, imputation and quality control

DNA was extracted from whole blood using a manual (2) or a semi-automatic (Autopure, Qiagen, Hilden, Germany) salting out method.

Samples were genotyped using the Illumina Infinium PsychArray (Illumina San Diego, CA, USA). Pre-imputation quality control included filtering SNPs on genotype call rate <98%, HWE *P*<10^-6^, and MAF = 0, and individuals on sample call rate <99%, relatedness and gender mismatch. Pre-phasing and imputation to the 1000 Genomes phase 3 reference panel was performed using IMPUTE2 version 2.3.2 (3). After imputation, genotype probabilities were converted to hard-called genotypes in Plink using a cut-off of 90%, meaning that only genotypes with a probability of more than 90% were called. Variants with no genotype probabilities above 90% were set to missing. Post-imputation quality control included filtering variants on MAF < 0.01 and INFO < 0.30, and the removal of CNVs and duplicate variants.

###### Analysis

Analyses were performed using the supplied scripts including age at phenotyping, gender and 20 principal components as covariates.

###### Acknowledgements

This study was supported by The National Program for Research Infrastructure 2007 (grant no. 09-063256), the Danish Agency for Science Technology and Innovation, and the US National Institute of Health (P01 AG08761). Genotyping was supported by NIH R01 AG037985 (Pedersen). Genotyping was conducted by the SNP&SEQ Technology Platform, Science for Life Laboratory, Uppsala, Sweden (http://snpseq.medsci.uu.se/genotyping/snp-services/).

##### DiscovEHR Study

###### Overview

Geisinger is the largest health care provider in central Pennsylvania with ~2 million patients. The DiscovEHR study, a subset of Geisinger’s MyCode Community Health Initiative cohort, with electronic medical records linked to genetic data (<http://www.discovehrshare.com/>). Geisinger patients enrolled in MyCode consent to broad research use of their samples. At the time of analysis, DiscovEHR included 92,476 consented individuals 92,476 with genetic data and with a median of 14 years of follow-up with EHR-derived clinical data.

For the purposes genetically-informed European-ancestry individuals were used for all analyses. For BMI and height adults aged 20-70 years. For height, unrealistic and outlier values were excluded (<44 inches, > 84 inches, and +/- 5 SD from mean). Height measurements were subsequently converted to metric (cm). Unrealistic and outlier weights <52 lbs, >650 lbs, and +/- 5 SD from mean were excluded. Additionally, weights during or within six months of pregnancy or following weight loss surgery were excluded. Weight in lbs was subsequently converted to metric units (kg). For individuals with repeated weight measurements. Cleaned height and weight were used to calculate BMI (kg/m^2) following data cleaning and conversion to metric units. For individuals with more than one measured height, median height in cm was used in all analyses. For repeated BMI, maximum BMI was used on all GWAS analyses. For SBP measures were extracted from clinical labs for adults aged 18 to 89 years. SBP measures were excluded on or following a diagnosis of chronic kidney disease (ICD-9 585), congestive heart failure (ICD-9 428), secondary hypertension (ICD-9 405). Additionally, SBP measurements were excluded if patient reported severe pain during encounter. SBP measurements were adjusted by adding 15 mmHg to measurement if patient was currently taking blood pressure lowering medication. For patients with multiple measures, median SBP was used in GWAS analyses.

###### Genotyping, imputation and quality control

Genotyping and quality control of DiscovEHR data have been previously described (Staples et al. 2018). Briefly, of the 92,476 with array-based genotyping, 67% were genotyped using the Illumina HumanOmniExpressExome (HOEE) genotyping platform with remainder typed on the Illumina Global Screening Array (GSA). These data are processed with Illumina’s GenomeStudio, imputed to the Haplotype Reference Consortium data and merged, resulting in 7.6 million variants. All array based data are QC’d with standard quality control procedures before association testing (Turner et al. 2011). Following imputation, variant dosages were filtered on INFO>0.3 and MAF>0.01, following the current study recommendations.

###### Analysis

All GWAS analyses were adjusted for sex, age (concurrent to phenotype measurement as described above, i.e. age at maximum BMI), array platform, and the first 20 PCs calculated from whole exome sequences.

##### Finnish Twin Cohort

###### Overview

The Finnish Twin Cohort consists of three longitudinal twin-family cohorts (1,2,3). The oldest cohort was initiated in 1975 and consists of twin pairs born before 1958 (3), while the FinnTwin12 (twins born 1983-1987) and FinnTwin16 (twins born 1975-1979) started in the 1990s (1,2). Each cohort has participated in 4 to 5 waves of data collection by mailed or online questionnaires, while some twin pairs and siblings have been invited to in-person studies, or telephone interviews with collection of DNA from venous blood samples or saliva. Phenotypes were compiled from these data sources such that the data from a twin pair was always at the same time point. The individual data collections have been approved by the respective data authorities, ethical committees and IRBs as documented in our reviews (1,2,3). Most sibling pairs included in the analyses are dizygotic twins, but some are also non-twin siblings.

###### Genotyping, imputation and quality control

Genotyping was done using Illumina Human610-Quad v1.0 B and Human670-QuadCustom v1.0 arrays at the Wellcome Trust Sanger Institute (Cambridge UK), Illumina HumanCoreExome- (12 v1.0 A, 12 v1.1 A, 24 v1.0 A, 24 v1.1 A, 24 v1.2 A) arrays at the Broad Institute of MIT and Harvard (MA, USA), Wellcome Trust Sanger Institute (Cambridge, UK), University of Chicago Genomics Facility (Chicago IL, USA) and Institute for Molecular Medicine Finland (Helsinki, Finland) and with Affymetrix FinnGen Axiom array at Thermo Fisher Scientific (Santa Clara CA, USA). The algorithm for genotype calling were Illumina’s GenCall for all HumanCoreExome chip genotypes, Illuminus for 610k & 670k chip genotypes and AxiomGT1 for Affymetrix chip genotypes. On Illumina arrays where genotypes were called to Illumina’s TOP strand, strand were flipped to forward strand using strand files generated by Will Rayner (<https://www.well.ox.ac.uk/~wrayner/strand/>). The genome build of all genotypes were set to GRCh37/hg19. In case where genotypes were called to NCBI36/hg18 or GRCh38/hg38, genome positions were lifted to GRCh37/hg19 using University of California Santa Cruz LiftOver program [4] with appropriate chain file. Genotype quality control were done in three batches (batch1: 610k+670k, batch2: HumanCoreExome and batch3: Affymetrix chip genotypes). Variants with call rate below 97.5% (batch1 and batch3) or 95% (batch2), samples with call rate below 98% (batch1) or 95% (batch2 and batch3), variants with minor allele frequency below 1% with Hardy-Weinberg Equilibrium p-value lower than 1e-06 were removed. In addition, amples from all batches with heterozygosity test method-of-moments F coefficient estimate value below -0.03 or higher than 0.05 (batch1 and batch2) or ±4SD from the mean (batch3) were removed along with the samples which failed sex check or were among the multi-dimensional scaling principal component analysis outliers. Total amount of genotyped autosomal variants after quality control were 475526 (batch1), 239894 (batch2) and 388673 (batch3) with following number of samples remaining for imputation: 2617 (batch1), 5328 (batch2) and 8218 (batch3). We then performed pre-phasing using Eagle v2.3 [5] and imputation with Minimac3 v2.0.1 using University of Michigan Imputation Server [6]. Genotypes of all batches were imputed to Haplotype Reference Consortium release 1.1 reference panel [7]. The final study sample were extracted from the data where all batches of imputed data were merged. As post-imputation quality control variants with MAF below 1% and imputation quality score below 0.5 were removed.

###### Analysis

Analyses were performed with the scripts supplied with this project using age, sex and 20 principal components as covariates.

7. the Haplotype Reference Consortium. A reference panel of 64,976 haplotypes for genotype imputation. Nature Genetics 2016. PMID: 27548312

##### Generation Scotland

###### Overview

The Generation Scotland: Scottish Family Health Study (GS:SFHS) is a family-based population cohort with DNA, biological samples, socio-demographic, psychological and clinical data from approximately 24,000 adult volunteers across Scotland. Although data collection was cross-sectional, GS:SFHS became a prospective cohort due to of the ability to link to routine Electronic Health Record (EHR) data. Over 20,000 participants were selected for genotyping using a large genome-wide array.

###### Genotyping, imputation and quality control

GS_SFHS waas genotyped using either the HumanOmniExpressExome8v1-2_or the HumanOmniExpressExome-8v1_A . Pre-imputation quality control included filtering SNPs on genotype call rate <98%, HWE *P*<10^-6^, and individuals on sample call rate <98%. MAF was >0.01 for OMNI markers and >0.0001 for Exome Chip markers. 602450 SNPS were phased using Shapeit v2.r873 + duohmm and imputed imputed using the Haplotype Research Consortium (HRC.r1-1) dataset.

###### Analysis

Analyses were performed using the supplied scripts including age at phenotyping, gender and 20 principal components as covariates.

###### Acknowledgements

Generation Scotland received core support from the Chief Scientist Office of the Scottish
Government Health Directorates [CZD/16/6] and the Scottish Funding Council [HR03006] and
is currently supported by the Wellcome Trust [216767/Z/19/Z]. Genotyping of the GS:SFHS
samples was carried out by the Genetics Core Laboratory at the Edinburgh Clinical Research
Facility, University of Edinburgh, Scotland and was funded by the Medical Research Council
UK and the Wellcome Trust (Wellcome Trust Strategic Award “STratifying Resilience and Depression Longitudinally” (STRADL) Reference 104036/Z/14/Z. C.H is supported by an MRC University Unit Programme Grant MC_UU_00007/10 (QTL in Health and Disease).

We are grateful to all the families who took part, the general practitioners and the Scottish
School of Primary Care for their help in recruiting them, and the whole Generation Scotland
team, which includes interviewers, computer and laboratory technicians, clerical workers,
research scientists, volunteers, managers, receptionists, healthcare assistants and nurses

##### Genetic Epidemiology Network of Arteriopathy (GENOA)

###### Overview

The Genetic Epidemiology Network of Arteriopathy (GENOA) study is a community-based study of hypertensive sibships that was designed to investigate the genetics of hypertension and target organ damage in African Americans from Jackson, Mississippi and non-Hispanic whites from Rochester, Minnesota (Daniels, 2004). In the initial phase of GENOA (Phase I: 1996-2001), all members of sibships containing ≥ 2 individuals with essential hypertension clinically diagnosed before age 60 were invited to participate, including both hypertensive and normotensive siblings. Exclusion criteria of the GENOA study were secondary hypertension, alcoholism or drug abuse, pregnancy, insulin-dependent diabetes mellitus, or active malignancy. Eighty percent of African Americans (1,482 subjects) and 75% of non-Hispanic whites (1,213 subjects) from the initial study population returned for the second examination (Phase II: 2001-2005). Study visits were made in the morning after an overnight fast of at least eight hours. Demographic information, medical history, clinical characteristics, lifestyle factors, and blood samples were collected in each phase. Written informed consent was obtained from all subjects and approval was granted by participating institutional review boards. Phenotypes and covariates were first processed in SAS 9.4.

All phenotypes were from GENOA Phase I except the cognitive phenotypes which were taken from an ancillary study conducted approximately 1 year after Phase II. Height in centimetres, BMI, and waist-to-hip ratio were calculated from measurements taken at in-person examinations. WHR was then adjusted for BMI with the residuals from linear regression used for this analysis. Education was the number of years of schooling. Sitting systolic blood pressure (SBP) (mmHg) was measured three times with a random zero sphygmomanometer. The average of the last two measurements was used in this study and adjusted by 10 units if an antihypertensive medication was taken. The general cognitive function score was the first unrotated principal component constructed by Principal Component Analysis (PCA) using the following 5 test measures: 1) the REY Auditory Verbal Learning Delayed Recall Test, 2) the Digital Symbol Substitution Test, 3) the FAS Word Fluency Test, 4) the Stroop Color Word Test, and 5) Part A of the Trail Making Test. Smoking phenotypes were assessed from self-report and include average cigarettes per day, ever smoker (more than 100 lifetime cigarettes), and age of initiation (>30 set to missing). Alcohol consumption was the average number of alcoholic units consumed per week as reported by the participant, aggregated across all types of alcohol, and outliers >5SD from the mean were removed. CRP and cholesterol measures were measured from blood. CRP was natural log transformed and standardized to mean 0 and SD 1. LDL was calculated as (total cholesterol) - HDL - (triglycerides/5), and all cholesterol measures were standardized to mean 0 and SD 1. eGFR was calculated from the CKD-EPI equation. Physical activity was defined as more than 20 minutes per week of moderate/vigorous activity.

###### Genotyping, imputation and quality control

All GENOA participants were genotyped on the Affymetrix Genome-Wide Human SNP Array 6.0 or the Illumina Human 1M-Duo BeadChip. Samples were removed if they had a missing call rate ≥0.05 or were an outlier ≥6 standard deviations from the mean of the first 10 genome-wide principal components from genotype data. SNPs were filtered to include those with MAF >0.01 and r 2 (INFO) >0.3. Imputation was done on the Michigan Imputation server (https://imputationserver.sph.umich.edu/index.html#!pages/home) using SHAPEIT and minimac3. Genotypes were imputed to the Haplotype Reference Consortium (HRC version r1.1) reference panel.

###### Acknowledgements

Support for the Genetic Epidemiology Network of Arteriopathy (GENOA) was provided by the
National Heart, Lung and Blood Institute (U01 HL054464, U01 HL054457, U01 HL054481, R01 HL087660, R01 HL085571, R01 HL119443) and the National Institute of Neurological Disorders and Stroke (R01 NS041558). Genotyping was performed at the University of Texas Health Sciences Center. We would like to thank the families that participated in the GENOA study.

##### HUNT

###### Overview

The Trøndelag Health Study (The HUNT Study) is a longitudinal population-based health study conducted in the county of Trøndelag, Norway. Data and samples have been collected through four surveys (HUNT1 [1984-1986], HUNT2 [1995-1997], HUNT3 [2006-2008] and HUNT4 [2017-2019]) (Krokstrand et al. 2013). At each time point, the entire adult population (≥ 20 years) was invited to participate by completing questionnaires, attending clinical examinations and interviews.

###### Genotyping, imputation and quality control

In total, DNA from 71,860 HUNT participants was genotyped using one of three different Illumina HumanCoreExome arrays (HumanCoreExome12 v1.0, HumanCoreExome12 v1.1 and UM HUNT Biobank v1.0). Quality control was performed at the marker and sample level. Samples that failed to reach a 99% call rate, had contamination > 2.5% as estimated with BAF Regress (Jun et al., 2012), large chromosomal copy number variants, lower call rate of a technical duplicate pair and twins, gonosomal constellations other than XX and XY, or whose inferred sex contradicted the reported gender, were excluded. Samples that passed quality control were analysed in a second round of genotype calling following the Genome Studio quality control protocol described elsewhere (Guo et al., 2014). Variants were excluded if (1) their probe sequences could not be perfectly mapped to the reference genome, cluster separation was < 0.3, Gentrain score was < 0.15, showed deviations from Hardy Weinberg equilibrium in unrelated samples of European ancestry with p-value < 0.0001, their call rate was < 99%, or another assay with higher call rate genotyped the same variant. Imputation was performed using Minimac3 (v2.0.1, <http://genome.sph.umich.edu/wiki/Minimac3>) (Das et al., 2016) and a merged reference panel that was constructed by combining the Haplotype Reference consortium release 1.1 (HRC v1.1) and a local reference panel based on 2,202 whole-genome sequenced HUNT study participants. SNPs were filtered to include those with MAF > 0.01 and Rsq > 0.3.

###### Analysis

Analyses were performed with the scripts supplied with this project using age, sex, 20 principal components and genotyping batch as covariates.

###### Acknowledgements

The HUNT Study is a collaboration between HUNT Research Centre, (Faculty of Medicine and Health Sciences, NTNU, Norwegian University of Science and Technology), Nord-Trøndelag County Council, Central Norway Regional Health Authority, and the Norwegian Institute of Public Health. The genotyping in HUNT was financed by the National Institutes of Health (NIH); University of Michigan; The Research Council of Norway; The Liaison Committee for Education, Research and Innovation in Central Norway; and the Joint Research Committee between St. Olavs hospital and the Faculty of Medicine and Health Sciences, NTNU. The genotype quality control and imputation has been conducted by the K.G. Jebsen Center for Genetic Epidemiology, Department of Public Health and Nursing, Faculty of Medicine and Health Sciences, NTNU, Norwegian University of Science and Technology.

##### Netherlands Twin Registry (NTR)

###### Overview

The Netherlands Twin Register (NTR, Ligthart et al. 2019) is an ongoing research initiative. Twins and their family members (siblings, offspring, spouses) participate on a voluntary basis. As of 2019 over 250,000 twins and family members have registered with the NTR, and over 26,000 participants have provided DNA samples for genome-wide SNP data. All adult participants have provided written informed consent, parents or primary caretakers have provided written informed consent for children. All data-collection protocols of the NTR are approved by the Medical Research Ethics Committee of the Amsterdam University Medical Centre and/ or the Ethical Review Board (VCWE) of the faculty of Behavior and Movement Science, Vrije Universiteit Amsterdam.

###### Genotyping, imputation and quality control

NTR participants were genotyped on the Affymetrix 6.0, Affymetrix Perlegen, Illumina Genomic Screening Array, Illumina Human660, or the Illumina Omni 1M array. Prior to imputation samples with a call rate below 90%, a mismatch between reported sex and biological sex, or an abnormal inbreeding F value (<-0.10 or >0.10) were removed. SNPs with a minor allele frequency (MAF) below 0.01, that deviate from Hardy-Weinberg Equilibrium (*p* < 1*10^-5^), have a missing rate of > 5%, or have >20 Mendelian errors were removed. These quality control steps were performed on each of the genotyping platforms independently. After this initial round of quality control SNPs were imputed to the Genome of the Netherlands (GONL, The Genome of the Netherlands Consortium, 2014). The resulting genetic data was filtered based on the same conditions mentioned earlier, and an additional filter to drop SNPs with a low imputation quality (*R^2^* < 0.90). The resulting dataset was then imputed a second time using the Haplotype Reference Consortium (HRC) version 1.1. After a final round of quality control, genetic principal components were generated from this dataset, and ethnic outliers based on these components were removed.

###### Analysis

Siblings were identified using genetic relatedness to ensure only full biological siblings were include. Both an additive genetic, and dominance genetic relatedness matrix were computed for all participants. Pairs of participants with an additive genetic relatedness between 0.475 and 0.625, as well as a dominance genetic relatedness between 0.125 and 0.375 were identified as biological siblings. For all phenotypes, except height and BMI, where multiple observations were available the last available observation was used. For height and BMI a direct measure was preferred over self-report, and the last available self-report was only used if no direct measure was available. If multiple direct measures of height or BMI were available, the last direct measure was used. Genetic analyses were performed using the scripts provided, with minor changes to fit local data storage formats. Age at survey completion, or measurement, of the included observation, sex, genotype platform and the first 20 principal components were used as covariates.

###### Acknowledgements

We would like to thank all members of twin families registered with the Netherlands Twin Register for their continued support of scientific research. NTR acknowledges past funding from the Netherlands Organization for Scientific Research (NWO) and The Netherlands Organisation for Health Research and Development (ZonMW) grants 904-61-090, 985-10-002, 912-10-020, 904-61-193,480-04-004, 463-06-001, 451-04-034, 400-05-717, Addiction-31160008, 016-115-035, 481-08-011, 400-07-080, 056-32-010, Middelgroot-911-09-032, OCW_NWO Gravity program –024.001.003, NWO-Groot 480-15-001/674, Center for Medical Systems Biology (CSMB, NWO Genomics), NBIC/BioAssist/RK(2008.024), Biobanking and Biomolecular Resources Research Infrastructure (BBMRI –NL, 184.021.007 and 184.033.111), X-Omics 184-034-019; Spinozapremie (NWO- 56-464-14192), KNAW Academy Professor Award (PAH/6635) and University Research Fellow grant (URF) to DIB; Amsterdam Public Health research institute (former EMGO+) , Neuroscience Amsterdam research institute (former NCA) ; the European Community's Fifth and Seventh Framework Program (FP5- LIFE QUALITY-CT-2002-2006, FP7- HEALTH-F4-2007-2013, grant 01254: GenomEUtwin, grant 01413: ENGAGE and grant 602768: ACTION); the European Research Council (ERC Starting 284167, ERC Consolidator 771057, ERC Advanced 230374), Rutgers University Cell and DNA Repository (NIMH U24 MH068457-06), the National Institutes of Health (NIH, R01D0042157-01A1, R01MH58799-03, MH081802, DA018673, R01 DK092127-04, Grand Opportunity grants 1RC2 MH089951, and 1RC2 MH089995); the Avera Institute for Human Genetics, Sioux Falls, South Dakota (USA). Part of the genotyping and analyses were funded by the Genetic Association Information Network (GAIN) of the Foundation for the National Institutes of Health. Computing was supported by NWO through grant 2018/EW/00408559, BiG Grid, the Dutch e-Science Grid and SURFSARA.

##### Orkney Complex Disease Study (ORCADES)

###### Overview

The Orkney Complex Disease Study (ORCADES) is a family-based, cross-sectional study that seeks to identify genetic factors influencing cardiovascular and other disease risk in the isolated archipelago of the Orkney Isles in northern Scotland (McQuillan et al., 2008). Genetic diversity in this population is decreased compared to Mainland Scotland, consistent with the high levels of endogamy historically. 2078 participants aged 16-100 years were recruited between 2005 and 2011, most having three or four grandparents from Orkney, the remainder with two Orcadian grandparents. Fasting blood samples were collected and many health-related phenotypes and environmental exposures were measured in each individual. All participants gave written informed consent and the study was approved by Research Ethics Committees in Orkney and Aberdeen (North of Scotland REC).

###### Acknowledgements

The Orkney Complex Disease Study (ORCADES) was supported by the Chief Scientist Office of the Scottish Government (CZB/4/276, CZB/4/710), a Royal Society URF to J.F.W., the MRC Human Genetics Unit quinquennial programme “QTL in Health and Disease”, Arthritis Research UK and the European Union framework program 6 EUROSPAN project (contract no. LSHG-CT-2006-018947). DNA extractions were performed at the Edinburgh Clinical Research Facility, University of Edinburgh. We would like to acknowledge the invaluable contributions of the research nurses in Orkney, the administrative team in Edinburgh and the people of Orkney.

##### QIMR Berghofer Medical Research Institute (QIMR)

###### Overview

Phenotypic data were collected during a series of longitudinal studies of Australian twins and their families. The content and details of data collection have been previously described 1–4. These studies were approved by the QIMR Berghofer Medical Research Institute Human Research Ethics Committee and the storage of the data follows national regulations regarding personal data protection. All participants provided informed consent.

###### Genotyping, imputation and quality control

Samples were genotyped using multiple Illumina HapMap, Omni, and GSA arrays. Pre-imputation quality control included excluding SNPs based on genotype call rate <95%, HWE P<10-6, MAF = 0, GenTrain Score < 0.6 [if data available], Mean GenCall Score < 0.7 [older Illumina array families only], unique position and strand alignment in a BLAST search, and sex-chromosome specific filters (female genotyping rate >1% for chr Y (not relevant); male heterozygosity >1% for chr X), and individuals based on sample call rate <99% and gender mismatch. Pre-phasing and imputation to the 1000G Phase3 v5 reference panel was performed using Eagle v2.4 (phased output) on the Michigan server. After imputation, SNPs were excluded based on MAF <0.001 and a minimum MAC=5. Genetic ancestry outliers were also excluded.

###### Analysis

Analyses were performed with the scripts supplied with this project using age, sex and 20 principal components as covariates.

###### Acknowledgements

We greatly thank the twins and their families for their participation. Thanks also to Grant Montgomery and his team for DNA collection and processing and to Scott Gordon for quality control and imputation of the genomic data. Data collection in the Australian sample was made possible by multiple grants from National Health and Medical Research Council and the Australian Research Council.

##### Swedish Twin Registry – Aging Samples (1 and 2)

###### Overview

The Swedish Twin Registry (STR) Aging samples included twins from five STR-based sub-studies: TwinGene (Zagai, Lichtenstein, Pedersen, & Magnusson, 2019), the Swedish Adoption Twin Study of Aging (SATSA) (Finkel & Pedersen, 2004), Aging in Women and Men (GENDER) (Gold, Malmberg, McClearn, Pedersen, & Berg, 2002), and Origins of Variance in the Oldest Old: Octogenarian Twins (OCTO-Twin) (McClearn et al., 1997) and the Study of Dementia in Swedish Twins (HARMONY) (Gatz et al., 2005).

Aging Sample I included 2764 dizygotic twins from the same pairs. TwinGene was collected between 2004 and 2008 with a total of 12 591 individuals. Phenotypic data, including Coronary heart disease (CHD), Type 2 Diabetes (T2D) and stroke, were collected by self-reported questionnaire. In addition, diagnostic coding (ICD10) for coronary heart disease, diabetes and stroke were obtained from the Swedish National Patient Register up until 2010. Systolic blood pressures (SBP) were measured in mm Hg. Blood samples of 50 ml was drawn from each individual by venipuncture at their local health-care facility and sent overnight to Karolinska University Laboratory. Low- and high-density lipoprotein (LDL and HDL, respectively), triglycerides (TG) and C-reactive protein (CRP) were measured by routine methods on semiautomated biochemistry analyzer (Beckman Coulter, CA). Hemoglobin A1c (HbA1C) was measured by high-liquid performance chromatography separation technique.

Aging Sample II included 850 dizygotic twins, including those from same-sex or opposite-sex pairs, with available genotyping on the Illumina Infinium Psych Array (Illumina San Diego, CA, USA), including SATSA, GENDER, Octo-Twin, and Harmony. Phenotypic data on height and BMI prioritized first available in-person measurements over self-reported values in the home study. Assayed triglycerides using standard methodswere taken from the first available in-person measurements in the home study, and per protocol values were standardized to mean zero and standard deviation one. Educational attainment, ever smoker, and depressive symptoms were collected via first available surveys in the home study. Education was coded into ISCED units and then to year equivalents per protocol. Depressive symptoms were measured from the 20-item Center for Epidemiologic Studies Depression Scale (CES-D), and per protocol standardized to mean zero and standard deviation one. Covariates included sex and age when all particpants were still alive (i.e., age in 1992).

Informed consents were obtained from all participants. The different study collections were approved separately by the regional ethical review board in Stockholm.

###### Genotyping, imputation and quality control

DNA was extracted from whole blood. For Aging Sample I, the serum was stored in liquid nitrogen. One 7ml EDTA tube of whole blood was stored in -80°C while a second 7ml EDTA tube of blood was used for DNA extraction. For both Aging Samples I and II, a 7ml EDTA tube of blood was used for DNA extraction using the Puregene extraction kit (Gentra systems, Minneapolis, USA). After extraction, the DNA was subsequently stored at -20°C.

Aging Sample I was genotyped on the Illumina OmniExpress bead chip array (700K). The quality controls included: individual missingness ≤0.03, genotype missingness ≤0.03, minor allele frequency ≥0.01, Hardy-Weinberg equilibrium P≥107, no sex mismatch, no excess heterozygosity (individuals with an F-statistic beyond 5 SDs from the sample mean), and no cryptic (unknown) relatedness. The genotyped data were imputed to The Haplotype Reference Consortium (HRC). Samples from the Aging sample II were genotyped on the Illumina Infinium Psych Array (Illumina San Diego, CA, USA). Quality control steps included: excluding markers not mapped to a chromosome, MAF = 0, not meeting Hardy-Weinberg equilibrium (p < 1e-6), and sex or relatedness discrepancies. After removing ambiguous strand SNPs, samples were pre-phased using SHAPEIT v2.r837 and imputed to 1000 genomes phase 1 version 3 using IMPUTE2 version 2.3.2 with default parameters.

###### Analysis

Phenotypic data preparation was performed in R-3.6.3 for Aging Sample I and SAS 9.4 (SAS, Inc., Cary, NC) for Aging Sample II. Preparation of genotype data and analyses were performed using the provided study scripts.

Aging Sample I - Analyses were performed using the provided scripts including age in 2019, sex and 20 principal components as covariates.

Aging Sample II - Analyses were performed using the provided scripts including age when all participants were still alive (i.e., age in 1992), sex and 10 principal components as covariates.

###### Acknowledgements

For STR Aging Sample II, PsychChip GWAS genotyping and collaborative work supported in part by the National Institutes of Health/National Institute on Aging grants R01 AG037985, R01 AG059329, and R01 AG060470 and DNA extraction by grants R01 AG17561 and R01 AG028555. Harmony was supported by grant R01 AG08724. OCTO-Twin was supported by grant R01 AG08861. Gender was supported by the MacArthur Foundation Research Network on Successful Aging, The Axel and Margaret Axson Johnson’s Foundation, The Swedish Council for Social Research, and the Swedish Foundation for Health Care Sciences and Allergy Research.SATSA was supported by grants R01 AG04563, R01 AG10175, the John D. and Catherine T. MacArthur Foundation Research Network on Successful Aging, the Swedish Council For Working Life and Social Research (FAS) (97:0147:1B, 2009-0795) and Swedish Research Council (825-2007-7460, 825-2009-6141).

We acknowledge The Swedish Twin Registry for access to data. The Swedish Twin Registry is managed by Karolinska Institutet and receives funding through the Swedish Research Council under the grant no 2017-00641. The content of this manuscript is solely the responsibility of the authors and does not necessarily represent the official views of the National Institutes of Health/National Institute on Aging.

10.1017/thg.2019.99

##### The Twins Early Development Study (TEDS)

###### Overview

The Twins Early Development Study (TEDS) is a multivariate, longitudinal study of >16,000 twin pairs representative of England and Wales, recruited 1994–1996 (Rimfeld et al. 2019). The demographic characteristics of TEDS participants and their families closely match those of families in the UK. Written informed consent was obtained from parents prior to data collection and from TEDS participants themselves past the age of 18. Current analyses were conducted on a sub sample of dizygotic (DZ) twin pairs with genome-wide genotyping and phenotypic data.

###### Genotyping, imputation and quality control

Two different genotyping platforms were used because genotyping was undertaken in two separate waves. AffymetrixGeneChip 6.0 SNP arrays were used to genotype 3,665 individuals. Additionally, 8,122 individuals (including 3,607 DZ co-twin samples) were genotyped on Illumina HumanOmniExpressExome-8v1.2 arrays. After quality control, 635,269 SNPs remained for AffymetrixGeneChip 6.0 genotypes, and 559,772 SNPs for HumanOmniExpressExome genotypes.

Genotypes from the two platforms were separately phased and imputed into the Haplotype Reference Consortium (release 1.1) through the Sanger Imputation Service before merging. Genotypes from a total of 10,346 samples (including 3,320 DZ twin pairs and 7,026 unrelated individuals) passed quality control, including 3,057 individuals genotyped on Affymetrix and 7,289 individuals genotyped on Illumina. The identity-by-descent (IBD) between individuals was < 0.05 for 99.5% in the merged sample excluding the DZ co-twins (range = 0.00 – 0.12) and ranged between 0.36 and 0.62 for the DZ twin pairs (mean = 0.49). There were 7,363,646 genotyped or well-imputed SNPs (for full genotype processing and quality control details, see (Selzam et al. 2018)).

For the current analyses, we further restricted to variants with high confidence imputation accuracy (INFO score >.75) and minor allele frequency >1%. The final number of SNPs was 6,300,709.

###### Analysis

Current analyses were adjusted for sample plate and chip, in addition to 20 principal components. Principal components were derived from a subset of 39,353 common (MAF > 5%), perfectly imputed (INFO = 1) autosomal SNPs, after stringent pruning to remove markers in linkage disequilibrium (r2 > 0.1) and excluding high linkage disequilibrium genomic regions to ensure that only genome-wide effects were detected.

###### Acknowledgements

We gratefully acknowledge the ongoing contribution of the participants in the Twins Early Development Study (TEDS) and their families. TEDS is supported by a program grant to RP from the UK Medical Research Council (MR/V012878/1 and previously MR/M021475/1), with additional support from the US National Institutes of Health (AG046938).

##### TwinsUK

###### Overview

TwinsUK is a cohort of volunteer adult twins from across the United Kingdom. The Registry was started in 1992 with the primary aim of assessment of heritability of osteoarthritis and osteoporosis in women. The success of early studies led to rapid evolution of the registry and it now incorporates about 12 000 twins, both male and female aged 18–103 years, studied for a whole range of clinical and behavioural traits.

###### Genotyping, imputation and quality control

5710 twins have undergone a genome-wide scan of either 317 000 single nucleotide polymorphism (SNP) markers (Illumina HumanHap300 Bead Chip) or 610, 000 SNPs (Illumina HumanHap610 Quad Chip). From these twins, 2840 participated in the first follow-up and 2545 in the second follow-up visits. The data was fully imputed using IMPUTE version 2 software, quality checked, and has been used in many international consortia for different phenotypes.

###### Acknowledgements

TwinsUK receives funding from the Wellcome Trust (212904/Z/18/Z) and European Union (H2020 contract #733100). TwinsUK and M.M. are supported by the National Institute for Health Research (NIHR)-funded BioResource, Clinical Research Facility and Biomedical Research Centre based at Guy’s and St Thomas’ NHS Foundation Trust in partnership with King’s College London. P.C. is founded by the European Union (H2020 contract #733100)

##### UK Biobank

###### Overview

UK Biobank is a large-scale prospective cohort study including 503,325 individuals aged between 38-73 years, who were recruited between 2006 and 2010 from across the United Kingdom. For the purposes of this study, we used a subsample of 40,210 siblings from 19,523 families. Full-siblings were derived using UK Biobank provided estimates of pairwise identical by state (IBS) kinships (>0.5-21*IBS0, <0.7) and IBS0 (>0.001, <0.008), the proportion of unshared loci.

###### Genotyping, imputation and quality control

UK Biobank study participants (N= 488,377) were genotyped using the UK BiLEVE (N= 49,950) and the closely related UK Biobank Axiom™ Arrays (N= 438,427). Directly genotyped variants were pre-phased using SHAPEIT3 and imputed using Impute4 and the UK10K, Haplotype Reference Consortium and 1000 Genomes Phase 3 reference panels.

###### Analysis

Analyses were performed using the supplied scripts including birth year, sex and 20 principal components as covariates.

###### Acknowledgements

UK Biobank received ethical approval from the Research Ethics Committee (11/NW/0383). Access to UK Biobank data was granted as part of application 15825 (PI: Dr Philip Haycock).

##### Viking Health Study – Shetland (VIKING I)

###### Overview

The Viking Health Study - Shetland (VIKING) is a family-based, cross-sectional study that seeks to identify genetic factors influencing cardiovascular and other disease risk in the population isolate of the Shetland Isles in northern Scotland (Kerr et al., 2019). Genetic diversity in this population is decreased compared to Mainland Scotland, consistent with the high levels of endogamy historically. 2105 participants were recruited between 2013 and 2015, most having at least three grandparents from Shetland. Fasting blood samples were collected and many health-related phenotypes and environmental exposures were measured in each individual. All participants gave informed consent and the study was approved by the South East Scotland Research Ethics Committee.

###### Acknowledgements

The Viking Health Study – Shetland (VIKING) was supported by the MRC Human Genetics Unit quinquennial programme grant “QTL in Health and Disease”. DNA extractions and genotyping were performed at the Edinburgh Clinical Research Facility, University of Edinburgh. We would like to acknowledge the invaluable contributions of the research nurses in Shetland, the administrative team in Edinburgh and the people of Shetland.

#### Phenotype definitions

Phenotype definitions were suggested in the analysis plan, based on previous GWAS.

##### Height

The participants height while standing measured in centimetres. If height has been recorded with greater precision, round to the nearest centimetre. Where height data have been collected from multiple sources, preference is that measures from direct assessments of cohort participants are used above self-reported height.

The meta-analysis units for height are centimetres.

##### BMI

Please calculate BMI as weight in kilograms divided by standing height in metres squared (BMI = kg/m2). Where weight has been collected with greater precision than kilograms, please round to the nearest kilogram. Where height has been collected with greater precision than centimetres (0.01 metre), please round to the nearest centimetre. Where weight and height data have been collected from multiple sources, our preference is that measures from direct assessments of cohort participants are used above self-reported measures.

The meta-analysis units for BMI are kg/m^2^.

##### Waist-hip ratio (WHR), adjusted for BMI

Please calculate waist-hip ratio adjusted for BMI by residualizing using linear regression. Waist and hip measures should be rounded to the nearest centimetre. Where multiple waist and hip measures are available our preference is for direct or clinic measurements above self-report. Please ensure that the waist-hip measurements and BMI were made concurrently. If there are multiple measurements over time, please choose the measurement occasion with the least missing data across your cohort.

The meta-analysis used untransformed WHR GWAS data.

##### Educational attainment

For years of education, completed academic qualifications should be mapped to levels of the International Standard Classification of Education (ISCED) and converted to the corresponding completed years of education required for completion of the qualification in US years of schooling. If ISCED levels 5 and 6 cannot be distinguished, please set values for those with tertiary education as 20 years of schooling.

The meta-analysis used standardised measures of years in full-time education (SD = 1).

##### Systolic blood pressure (SBP)

The participants systolic blood pressure adjusted for treatment with anti-hypertensives. Please use the time point that has the least missing data across your sample. Please replace missing values with values from other measurement occasion if multiple time points are available. If multiple measurements of blood pressure were taken at each clinic visit, please use the average of the measurements. If some participants have been treated with treated with anti-hypertensive medication, please add 10mmHg to the blood pressure of individuals who are on treatment at the time of measurement.

The meta-analysis units for SBP are mmHg.

##### Cognitive ability (performance on cognitive tests)

A general cognitive function score will be created, where appropriate data are available in the sample, by Principal Component Analysis (PCA) using at least 3 cognitive tests that assess different cognitive domains. Only one score should be used from each cognitive test. The tests should not include the clinical cognitive assessments that are used as screening instruments for dementia (e.g., MMSE). The score to be created and used for the genetic analysis will be the first unrotated principal component. The general cognitive function score from the PCA (first unrotated principal component) should be saved as a standardized variable (mean = 0; SD = 1). The variable must be in the direction of higher positive scores indicating better general cognitive performance.

The meta-analysis used standardised measures of cognitive ability (SD = 1).

##### Smoking behaviour

For each smoking phenotype only include information on cigarette smoking behaviour. Do not include information about pipes, cigars, or other forms of tobacco use.

For ever smoker, a binary measure recording those who report ever being a regular smoker in their life, either current or former.

For smoking intensity, the average number of cigarettes smoked per day as a current or former smoker (set values for non-smokers to missing, not zero). If numbers of cigarettes per day have been recorded as categorical (e.g. 1-5 cigarettes per day), set value as the mid-point of the range (e.g. 2.5). Please remove extreme outliers (e.g. 5 S.D. away from the mean).

The meta-analysis used risk increase (ever smoking) and untransformed cigarettes per day (smoking intensity) data.

##### Alcohol consumption

The average number of alcoholic units consumed per week as reported by the participant, aggregated across all types of alcohol. If numbers of drinks have been recorded as categorical (e.g. 1-5 drinks per week), set value as the mid-point of the range (e.g. 2.5). Please remove extreme outliers (e.g. 5 S.D. greater than the mean).

The meta-analysis used untransformed units of alcohol consumption.

##### Depressive symptoms

Depression is a highly heterogeneous disorder with many clinical presentations. The diagnosis requires a distinct change of mood, characterized by sadness or irritability, accompanied by psychophysiological changes, such as disturbances in sleep, appetite, or sexual desire; constipation; loss of the ability to experience pleasure in work or with friends; crying; suicidal thoughts; and slowing of speech and action.

A great variety of rating scales have been developed to assess symptoms of depression. The scales differ in the number of items included and with respect to the types of symptoms assessed: mood symptoms (depressed mood and irritability), behavioural symptoms (suicide and anhedonia), somatic symptoms (appetite disturbance, sleep disturbance, low energy, and psychomotor retardation or agitation), cognitive symptoms (hopelessness and worthlessness), and concentration symptoms (poor concentration and decision-making). Frequently used scales include: the Hamilton Rating Scale for Depression (HRSD), the Beck Depression Inventory (BDI), the Center for Epidemiologic Studies Depression Scales (CES-D), the Quick Inventory of Depressive Symptoms (QIDS), and the Self-Rating Depression Scale (SDS). In MoBa (one of the largest cohorts) depressive symptoms were measured by the Short Mood and Feelings Questionnaire (SMFQ) in adolescents, and by the Hopkins Symptom Checklist (H-SCL) in adults.

If multiple rating scales of depressive symptoms are available, select the scale with most items (tapping most symptom domains). Please sum the items and treat the final summed variable as a continuous measure of depression symptoms. Please ensure that measures are coded so that higher values represent higher symptom levels. Please standardize the score to have mean zero standard deviation one.

The meta-analysis used standardised measures of depressive symptoms (SD = 1).

##### Subjective wellbeing

Reporting of subjective wellbeing is likely to be highly variable across studies. Our preference is for subjective wellbeing to have been measured using a battery of questions. If your study has done this, please sum responses and treat this final summed variable as a continuous measure of subjective wellbeing. If data on subjective wellbeing are only available from a single question reported on a Likert scale, please treat the response variable as continuous. Please ensure that measures are coded so that higher values represent higher levels of subjective wellbeing.

The meta-analysis used standardised measures of subjective wellbeing (SD = 1).

##### Neuroticism

Neuroticism is a fundamental domain of personality functioning and structure. The personality trait refers to a lack of emotional stability; stress vulnerability; the tendency to experience intense negative emotions, affects, and cognitions; and impulsive behaviours under emotional strain. Neuroticism can be validly measured from age 6-7 years. The Hierarchical Personality Inventory for Children is one of few instruments specifically designed to assess neuroticism in children. For adults, scales include the NEO-Personality Inventory, the Eysenck Personality Questionnaire, and various scales using Big-Five factor markers from the International Personality Item Pool (IPIP- including the 300-item representation of the Revised NEO Personality Inventory). Most modern neuroticism scales show strong overlap in item content (although they consist of different items), and the sum scores correlate highly across inventories. Scales usually agree on lower-order traits (e.g. anxiety-withdrawal, depression-unhappiness, vulnerability-stress reaction). However, there is less agreement whether aggression, impulsivity, inferiority, and dependency belong to the neuroticism domain.

Please sum the neuroticism items and treat the final summed variable as a continuous measure. The scale should be standardized with mean=0 and SD=1. Please ensure that higher values on the scale represent higher levels of neuroticism.

The meta-analysis used standardised measures of neuroticism (SD = 1).

##### Number of biological children

Restricted to study participants who have reached the end of their reproductive window, defined as over the age of 45 if female and over the age of 55 if male.

*Number (of children) ever born (NEB)*

NEB can be treated as a continuous measure that has been asked directly or can be imputed from several survey questions (such as pregnancy histories). A standard question within most surveys asks:

*How many children have you given birth to?*

Another variant is:

*How many children do you have?*

In most cases it is also possible to distinguish between biological (live born or stillborn), adopted or stepchildren. When this information is available, we will refer to **live born** **biological children**. In some surveys, the birth date of each child is asked, enabling a calculation of NEB. In many surveys, this question has been adapted for male and female subjects and in several cases only women have been asked. Sometimes, there is a filter in the survey, asking whether the respondent has any children. For twin studies please include a count of 2 for all live born biological children. Individuals are eligible if they meet the following criteria:

1. They were assessed for NEB at least at age 45 for women, age 55 for men;
2. Those who have both given birth to a child (parous) and those who have not (nulliparous);
3. All relevant covariates (year of birth) are available for the individual;
4. They were successfully genotyped genome-wide (recommended individual genotyping rate > 95%);
5. They passed the cohort-specific standard quality controls, e.g. excluding individuals who are genetic outliers in the cohort.

The meta-analysis used the untransformed number of biological children.

##### Age at first birth (AFB)

AFB can be treated as a continuous measure, which has generally been asked directly or can be imputed from several survey questions (such as date of birth respondent and date of birth of first child). The most common question is:

*How old were you when you had your first child?*

Another variant is:

*What is the date of birth of your first child?*

In the case of the latter, you can simply create a new AFB variable by subtracting the date of birth of the first child from the date of birth of the respondent. In many surveys, this question has been adapted for male and female subjects and in some cases only women have been asked. Individuals are eligible if they meet the following criteria:

a.     They were assessed for AFB and have given birth to a child (parous); both for females and for males.

b.     All relevant covariates (year of birth) are available for the individual;

c.      They were successfully genotyped genome-wide (recommended individual genotyping rate > 95%);

d.     They passed the cohort-specific standard quality controls, e.g., excluding individuals who are genetic outliers in the cohort.

The meta-analysis used untransformed age at first birth.

##### C-reactive protein (CRP)

Please measured serum CRP in mg/L by using standard laboratory techniques and transform the values by natural log. Please exclude individuals with auto-immune diseases, individuals taking immune-modulating agents (if this information was available), and individuals with CRP values 4 SD or more away from the mean from CRP analyses (i.e. set to missing). Please normalize to mean zero and SD one.

The meta-analysis used standardised measures of CRP (SD = 1).

##### HbA1c

Trait are “raw” untransformed HbA1c values in % of hemoglobin, without rank normalization. Set all individuals with Diabetes (T2D, T1D) to missing ('diagnosed', on diabetes treatment (oral and insulin) or fasting plasma glucose (FPG) >=7 mmol/L). If FPG is NOT available, set individuals with 2hr-hour glucose >= 11.1 mmol/L and/or HbA1c >= 6.5%. Set participants with major blood abnormalities (thalassemia, sickle cell anemia, etc) or those who have had a blood transfusion in the previous 2-3 months. Definition taken from Wheeler et al. (2017).

 The meta-analysis used standardised measures of HbA1c (SD = 1).

##### Lipids

The participants untreated LDL-cholesterol, HDL-cholesterol and triglyceride levels. Please replace missing values with values from other measurement occasion if multiple time points are available. If multiple measurements of cholesterol were taken at each clinic visit, please use the average of the measurements. Please normalise to mean zero standard deviation one.

The meta-analysis used standardised measures of LDL, HDL and TG (SD = 1).

##### Lung function (FEV1/FEV1FVC)

Restrict dataset to those individuals with no missing data for the ever smoking or never smoking variable and to those with complete data on both forced expiratory volume (FEV1) and forced vital capacity (FVC). Undertake linear regression of age, age squared, sex, height on FEV1 and use residuals for all subsequent analyses. Transformation would be taken once for each trait (FEV1 and FEV1/FVC ratio) and used for all analyses (including subgroups). Transform residuals to ranks and then to normally distributed z-scores. These inverse-normal transformed residuals are then used as the phenotype for association testing under an additive genetic model.

Never-smokers only: Repeat analysis as for 1

Ever-smokers only:  Repeat analysis as for 1

**Repeat** the above for outcomes: **FEV1/FVC ratio**

The meta-analysis used standardised measures of FEV1 and FEV1/FVC ratio (SD = 1).

##### Age at menarche

Age at menarche should be treated as a continuous measure in years, which has generally been asked directly of study participants. The most common question is:

*How old were you when you had your first menstrual period?*

In the event of responses being recorded in multi-year categories please set the response as the year midpoint (For example, the response category 8-9 would be recoded as 8.5). See Day et al (2017) for further information.

*Day et al. Genomic analyses identify hundreds of variants associated with age at menarche and support a role for puberty timing in cancer risk. 2017. Nature Genetics, 49:834–841.*

The meta-analysis used untransformed age at menopause.

##### Age at menopause

Age at menopause should be treated as a continuous measure in years, which has generally been asked directly of study participants. The most common question is:

*At what age did your natural periods cease?*

If the information is available, women who had radiation or chemotherapy or surgically induced menopause should be excluded from the analysis. In line with He et al (2007) we suggest that women who report ages at menopause earlier than age 40 or later than age 60 should be excluded.

*He et al. Genome-wide association studies identify loci associated with age at menarche and age at natural menopause. 2009. Nature Genetics, 41:724–728.*

The meta-analysis used untransformed age at menarche.

##### Estimated glomerular filtration rate (eGFR)

1) If necessary correct the serum creatine measurements. This will depend on the measurement method (example given for HUNT)

- In HUNT serum creatine was measured using the Isotope Dilution Mass Spectroscopy Method (IDMS).

a) For HUNT2 either use corrected serum creatine measurement or apply the correction formula (1.11 x s-creatinine HUNT2 -27.4).

b) For HUNT3, correct the serum creatine measurement using the formula (0.889 x s-creatinine HUNT3 +12.6) x 1.11 -27.4)

2) Use corrected serum creatine to calculate eGFR by CKD-EPI (4-equations) model. Here are the 4 equations:

a) For "Female" & SeCrea <=61.9: eGFR= (144 * (SeCrea/61.9)^-0.329 * (0.993)^age)

b) For "Female" & SeCrea2>61.9: eGFR= (144 * (SeCrea/61.9)^-1.209 * (0.993)^age)

c) For "Male" & SeCrea<=79.6: eGFR= (141 * (SeCrea/79.6)^-0.411 * (0.993)^age)

d) For "Male" & SeCrea>79.6: eGFR= (141 * (SeCrea/79.6)^-1.209 * (0.993)^age)

The meta-analysis used untransformed eGFR.

##### Physical activity

Moderate-to-vigorous intensity leisure time activity (more vs. less than 20 min/week)

Equal to or more than >  20 minutes of “strenuous sports” and “other exercise”  combined   (dichotomized)

Equal to or more than > 20 minutes of physical activity => METs 3   (dichotomized)

The meta-analysis used risk increase in the binary physical activity measure.

### Supplementary Figures

#### Supplementary Figure 1 Shrinkage for height across European ancestry cohorts

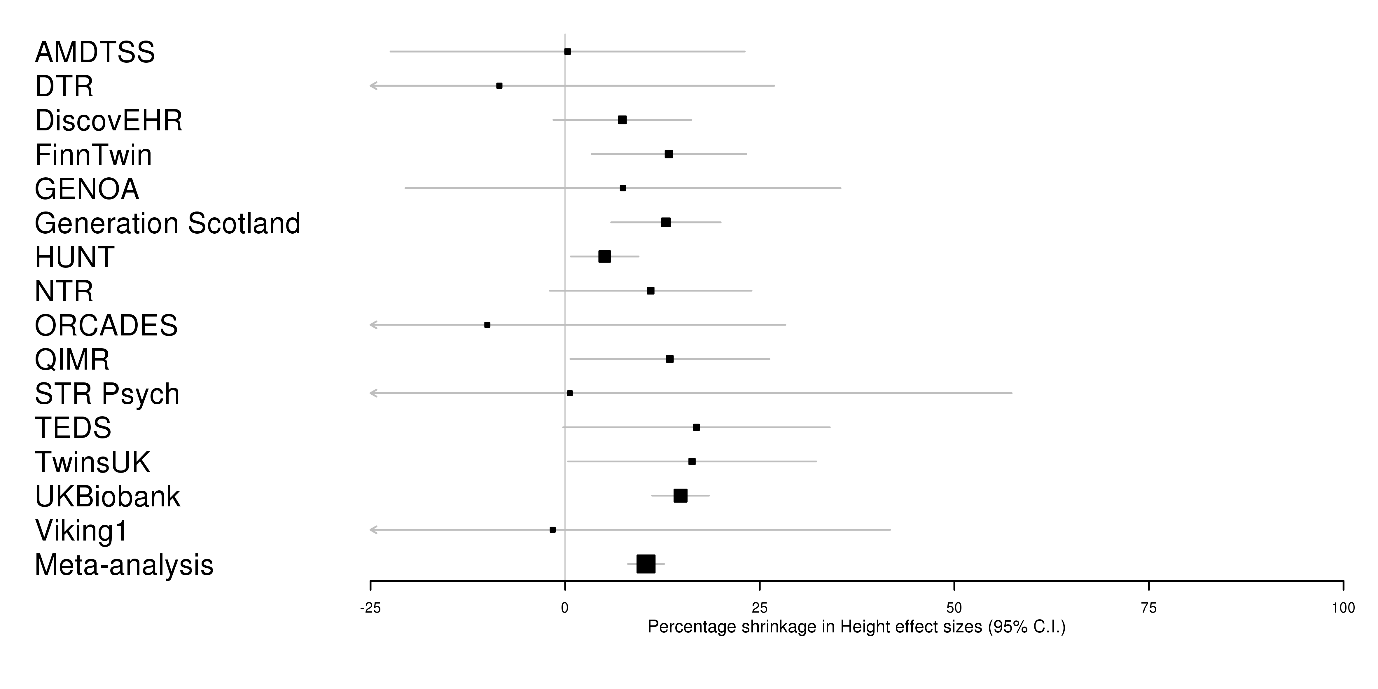

AMDTSS = Australian Mammographic Density Twin Study, DTR = Danish Twins Registry, NTR = Netherlands Twin Registry, QIMR = QIMR Berghofer Medical Research Institute (QIMR), TEDS = Twins Early Development Study.

Supplementary Figure 1 shows estimates of within-sibship shrinkage (% attenuation in effect sizes from population estimates to within-sibship estimates) for height variants in all of the cohorts contributing to the European meta-analysis as well as the meta-analysis GWAS. These estimates used the weighted score for each phenotype at the more liberal threshold (P < 1x10^-5^).

#### **Supplementary Figure 2** Shrinkage for educational attainment across European ancestry cohorts

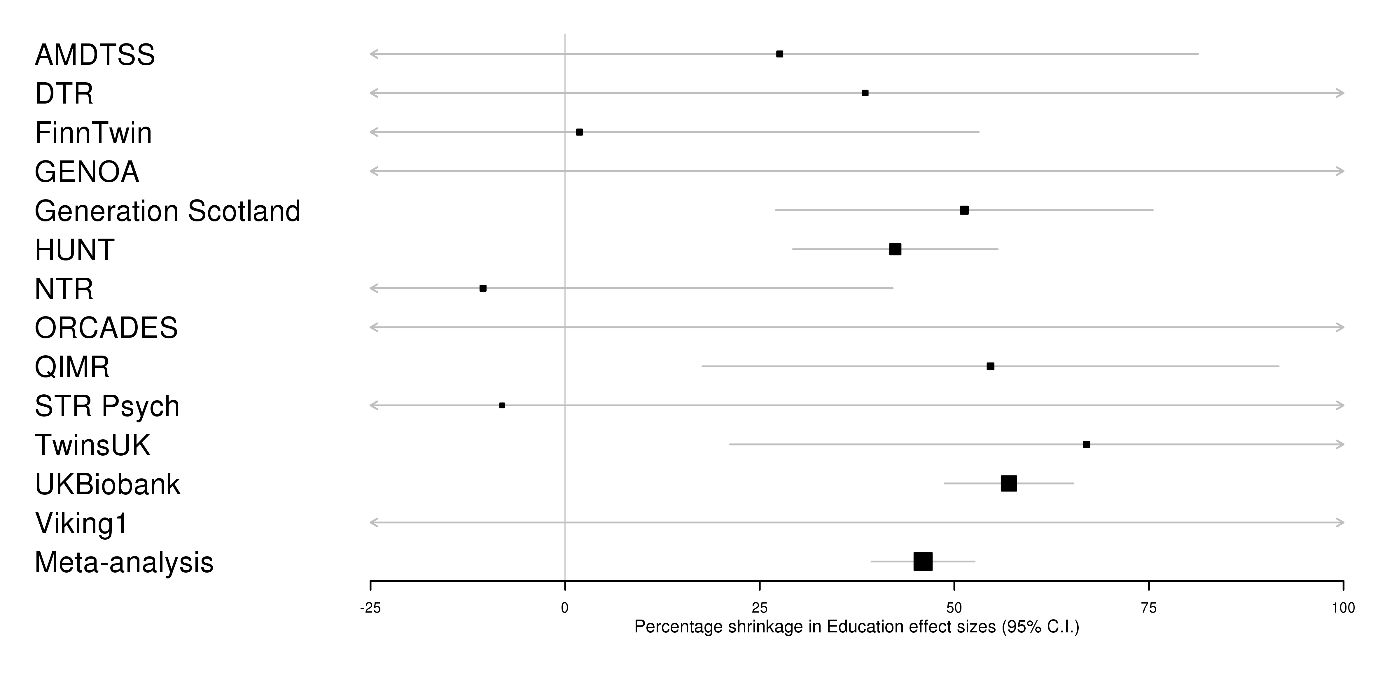

AMDTSS = Australian Mammographic Density Twin Study, DTR = Danish Twins Registry, NTR = Netherlands Twin Registry, QIMR = QIMR Berghofer Medical Research Institute (QIMR).

Supplementary Figure 2 shows estimates of within-sibship shrinkage (% attenuation in effect sizes from population estimates to within-sibship estimates) for educational attainment variants in all of the cohorts contributing to the European meta-analysis as well as the meta-analysis GWAS. These estimates used the weighted score for each phenotype at the more liberal threshold (P < 1x10^-5^).

#### Supplementary Figure 3 Shrinkage estimates from China Kadoorie Biobank

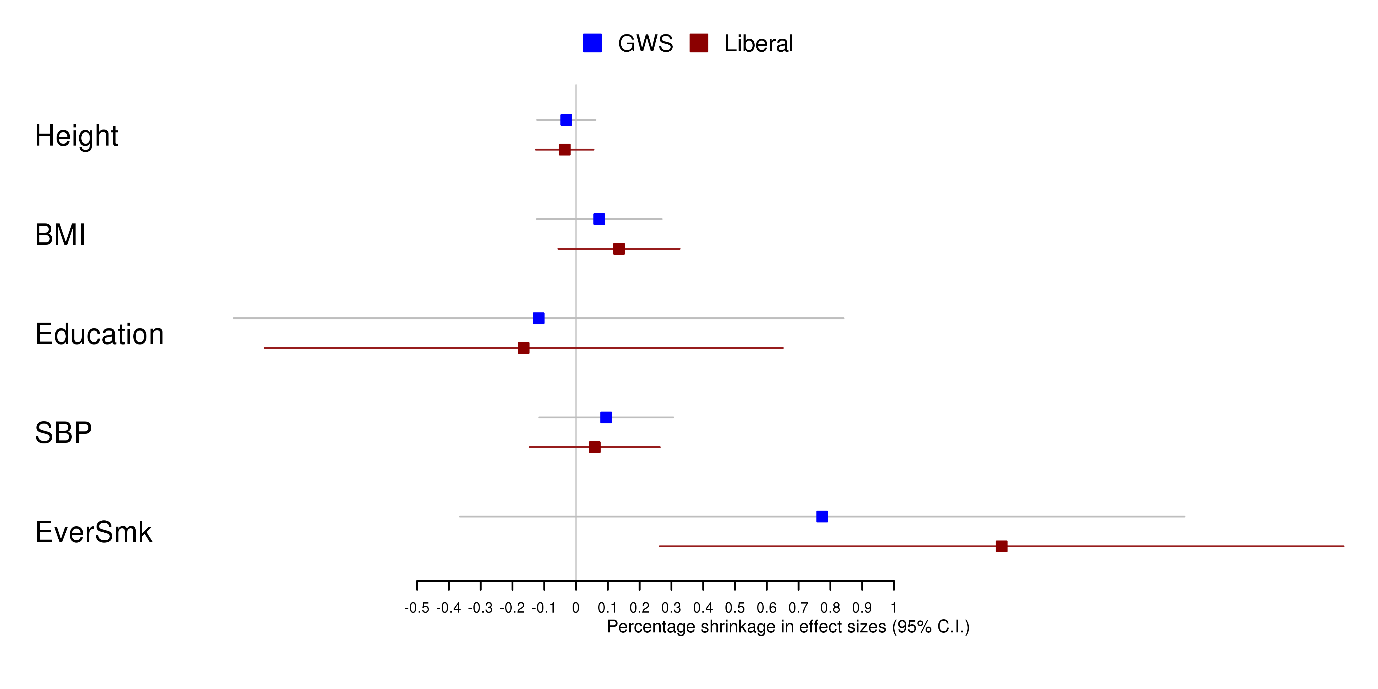

GWS = score including variants with P < 5x10^-8^, Liberal = score including variants with P < 1x10^-5^, BMI = body mass index, SBP = systolic blood pressure, EverSmk = ever smoking.

Supplementary Figure 3 contains within-sibship shrinkage (% attenuation in effect sizes from population estimates to within-sibship estimates) for height, BMI, educational attainment, systolic blood pressure and ever-smoking genetic variants in China Kadoorie Biobank. The figure includes genetic variants from the genome-wide significant (blue) and liberal (red) thresholds. Note that the genetic variants tested were identified in UK Biobank but any ancestral differences will likely equally affect both the population and within-sibship estimates, meaning that the shrinkage estimate will be unbiased.

#### Supplementary Figure 4 Evidence of polygenic adaption using SDS for 25 phenotypes.

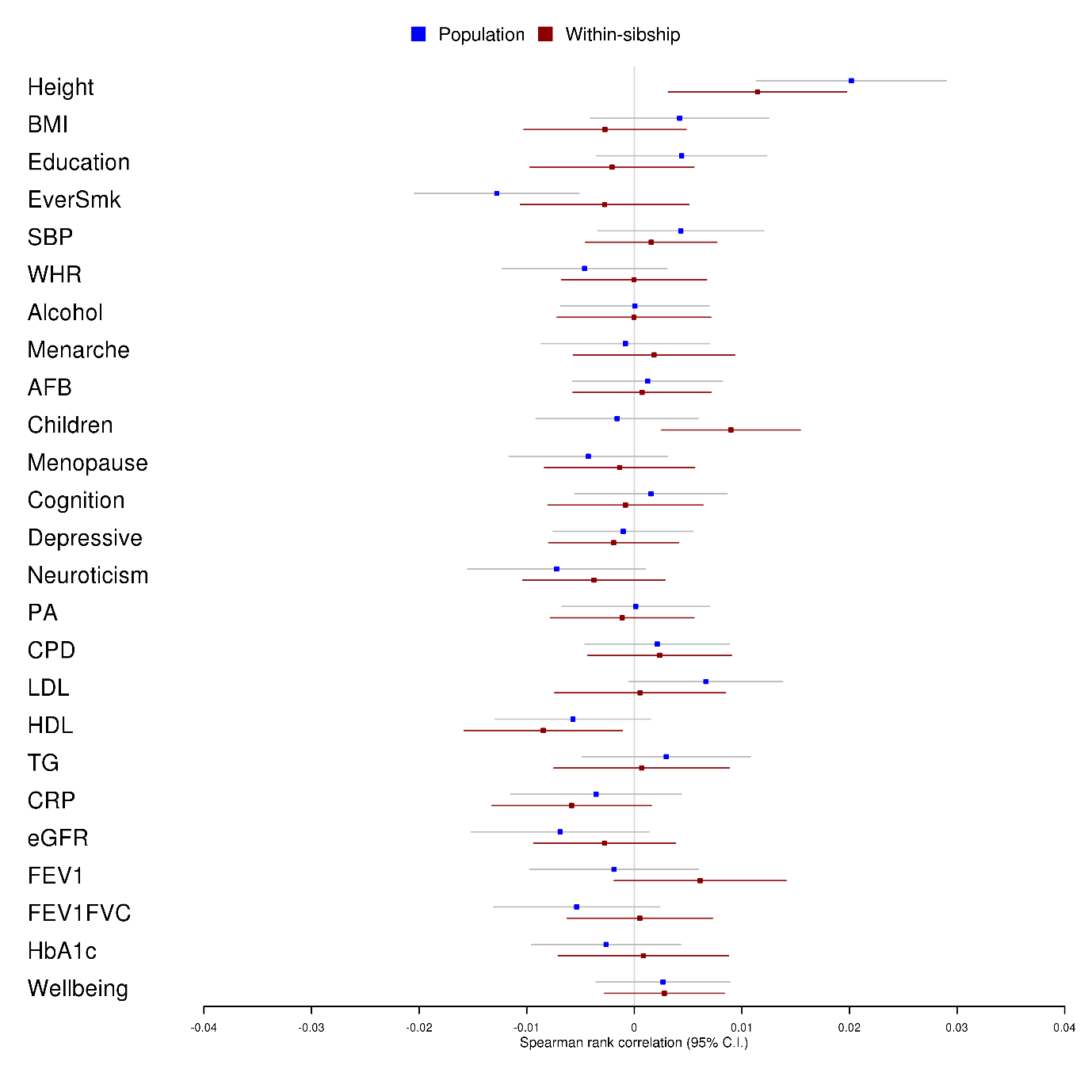

BMI = body mass index, EverSmk = ever smoking, SBP = systolic blood pressure, WHR = waist-hip ratio, AFB = age at first birth, PA = physical activity, CPD = cigarettes per day, TG = triglycerides, CRP = C-reactive protein, eGFR = estimated glomerular filtration rate, FEV1 = forced expiratory volume, FEV1FVC = ratio of FEV1/forced vital capacity, HbA1c = Haemoglobin A1C.

Supplementary Table 4 displays spearman rank correlation estimates between tSDS (SDS scores aligned with phenotype increasing alleles) and absolute phenotype Z scores for 25 phenotypes. The phenotype Z scores were taken from both the meta-analysis of population (blue) and within-sibship (red) estimates. Positive correlations indicate evidence of historical positive selection on phenotype increasing alleles.

#### Supplementary Figure 5 Scatter plot of SDS p-value bins against mean tSDS (of the bin) using the within-sibship height meta-analysis GWAS data.

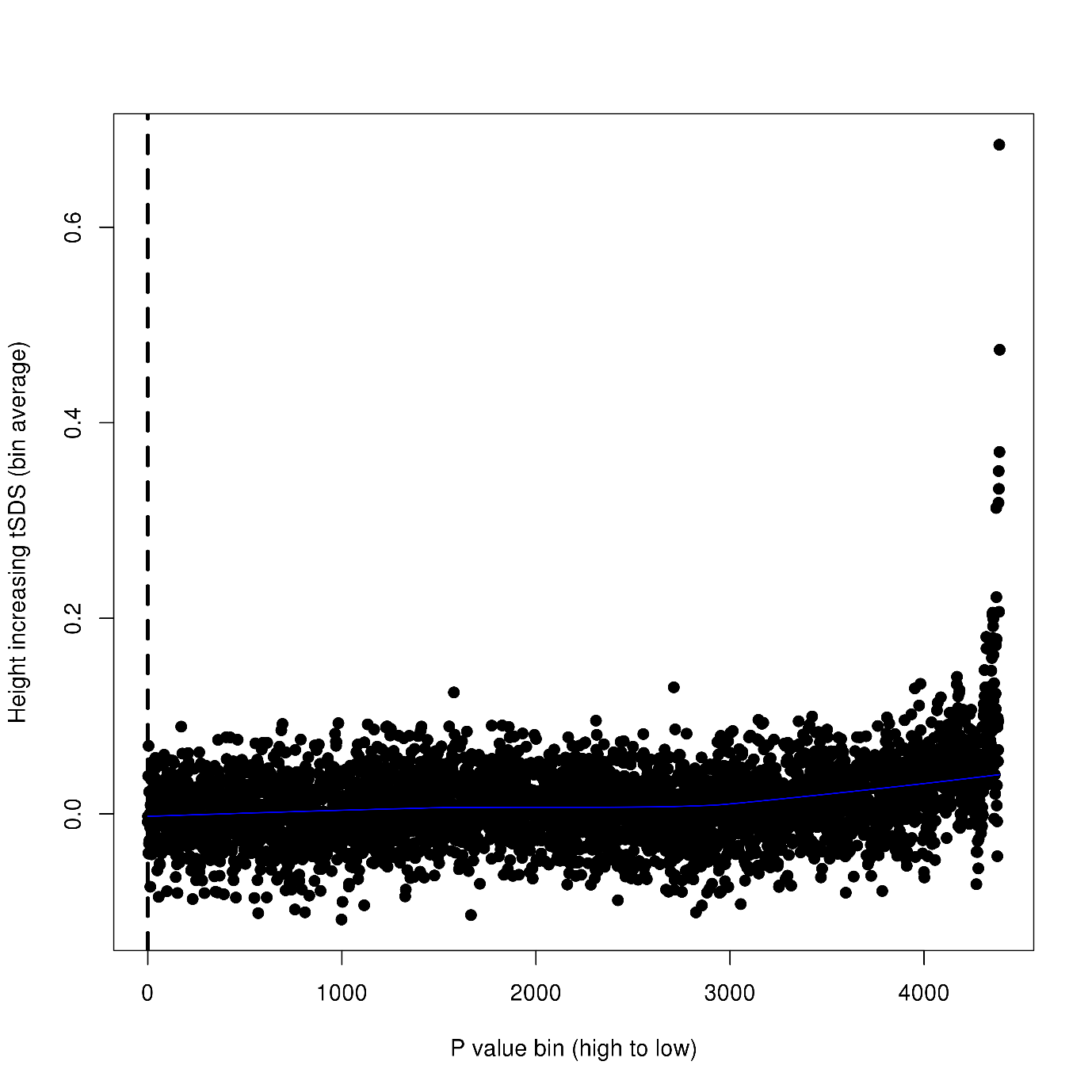

In Supplementary Figure 5 each data point is the mean tSDS (SDS score alligned with height increasing allele) in a set of 1000 genetic variants. Genetic variants were ordered by height P-value (from within-sibship meta-analysis GWAS data) and divided into bins. The plot illustrates evidence of a correlation between decreasing height P-value and higher mean tSDS suggestive of polygenic adaptation on height increasing alleles.

#### Supplementary Figure 6 Histogram of tSDS for independent variants associated with height in the within-sibship meta-analysis data (P < 1x10^-5^)

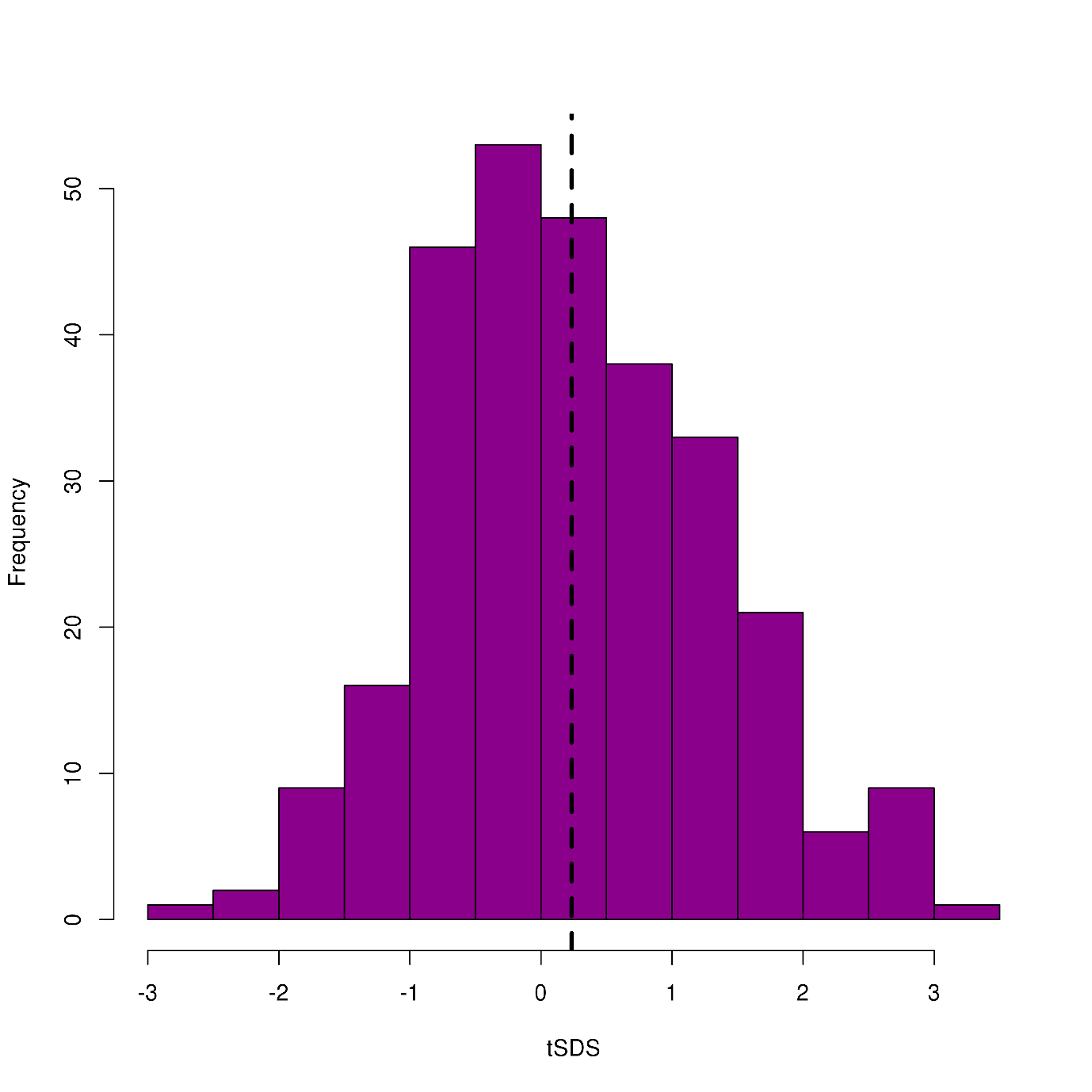

Supplementary Figure 6 is a histogram of the distribution of tSDS (SDS score aligned with height increasing alleles) amongst 310 putative independent height loci identified from the within-sibship meta-analysis data (P < 1x10^-5^). The plot indicates that the mean tSDS of these loci is higher than 0, consistent with polygenic adaptation on height increasing alleles.

#### Supplementary Figure 7 LDSC estimates of confounding across 25 phenotypes using within-sibship data.

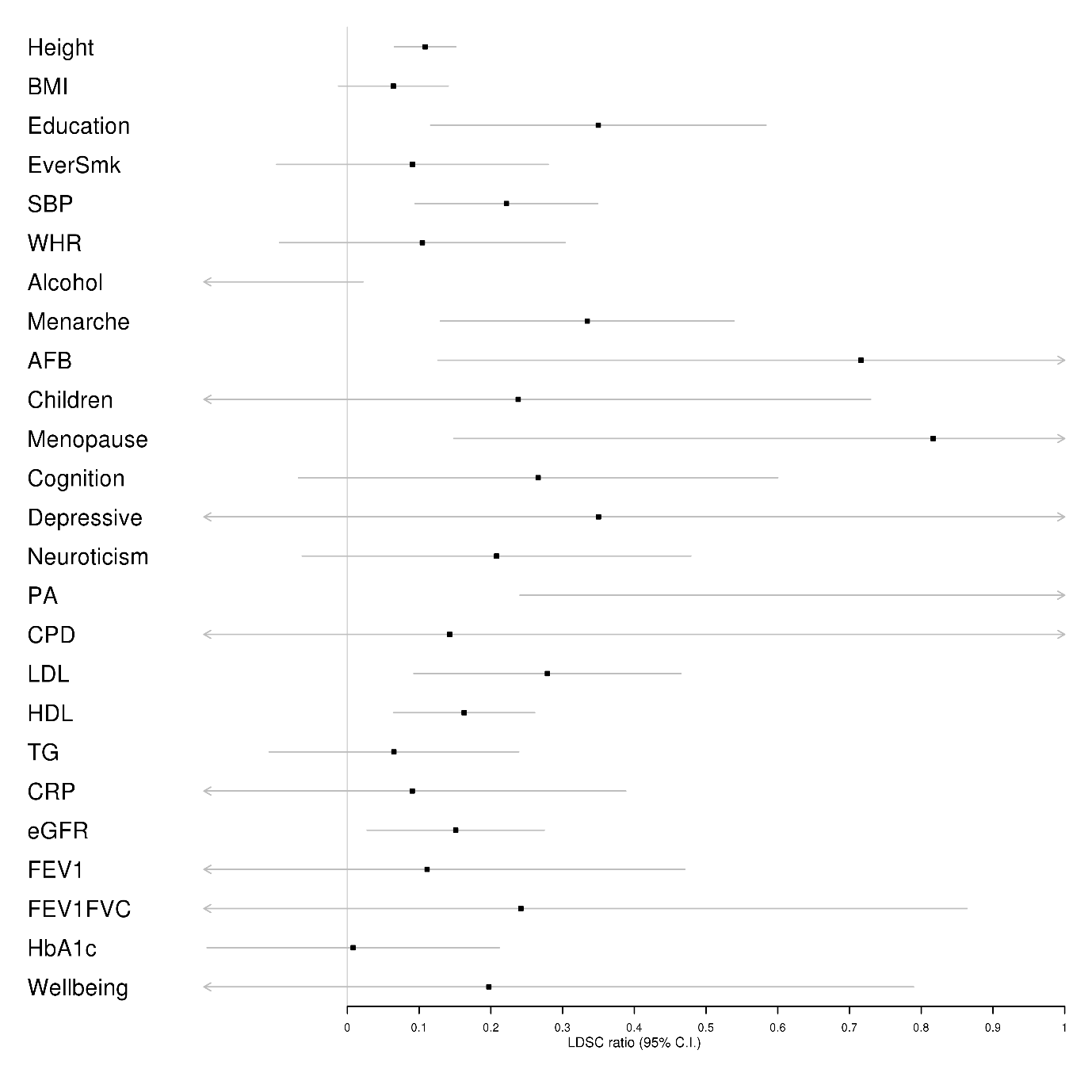

BMI = body mass index, EverSmk = ever smoking, SBP = systolic blood pressure, WHR = waist-hip ratio, AFB = age at first birth, PA = physical activity, CPD = cigarettes per day, TG = triglycerides, CRP = C-reactive protein, eGFR = estimated glomerular filtration rate, FEV1 = forced expiratory volume, FEV1FVC = ratio of FEV1/forced vital capacity, HbA1c = Haemoglobin A1C.

Supplementary Figure 7 shows LDSC ratio estimates (a measure of the % of the polygenic signal attributable to confounding) for 25 phenotypes using the within-sibship meta-analysis data.

#### Supplementary Figure 8 LDSC ratios from height GWAS

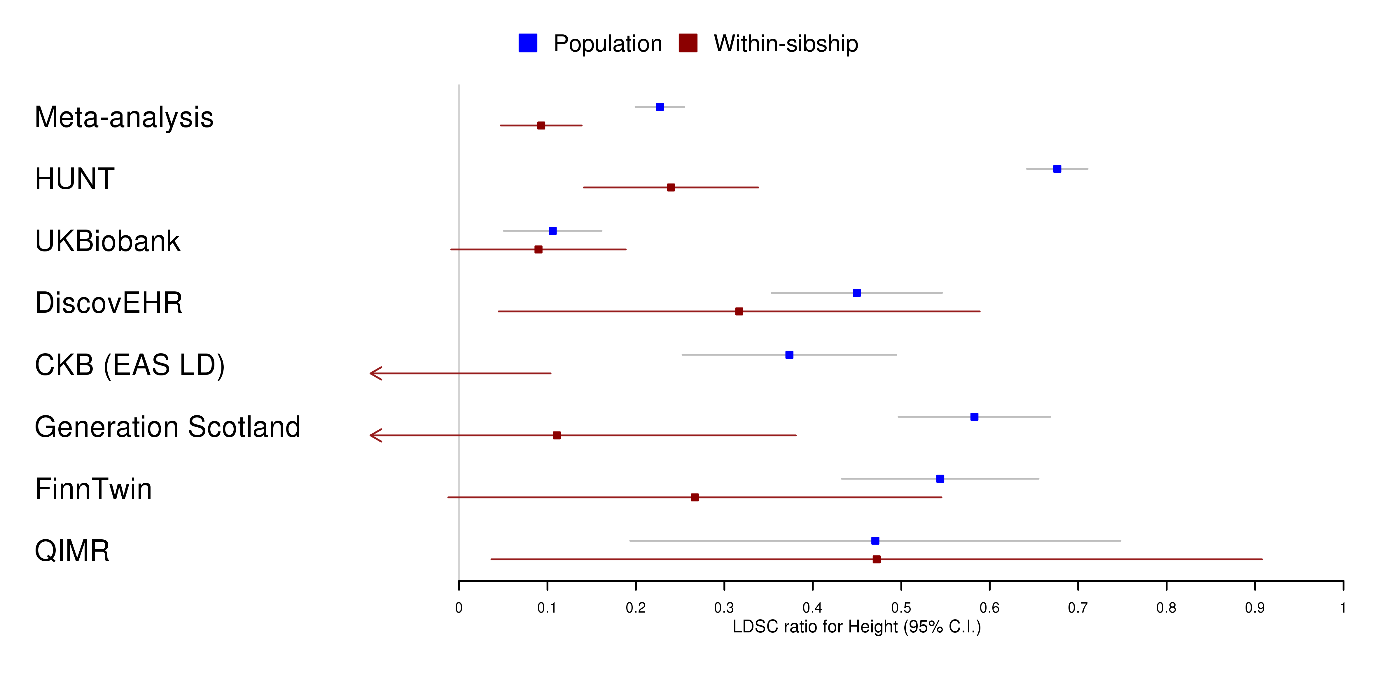

CKB = China Kadoorie Biobank, EAS LD = East Asian LD scores, QIMR = QIMR Berghofer Medical Research Institute.

Supplementary Figure 8 shows LDSC ratio estimates (a measure of the % of the polygenic signal attributable to confounding) for height from the summary data of 7 individual studies and the meta-analysis of European studies.
